# CILP Inhibits Hyaline Cartilage Fibrosis and Chondrocyte Ferroptosis via Keap1-Nrf2 Axis in Early Osteoarthritis Exercise Therapy

**DOI:** 10.1101/2024.11.02.621635

**Authors:** Shuangshuo Jia, Zhehan Hu, Zihan Li, Weiming Zhang, Liang Chen, Changping Niu, Ziqi Zhao, Yuhan Sun, Gang Yao, Yang Wang, Yue Yang

**Affiliations:** Department of Orthopedic Surgery, Shengjing Hospital of China Medical University, Shenyang, China; Shengjing Hospital of China Medical University, Department of Gastroenterology and Medical Research Center, Liaoning Key Laboratory of Research and Application of Animal Models for Environmental and Metabolic Diseases, ShenYang, Liaoning, 110000, China; Key Laboratory of Health Ministry for Congenital Malformation, Shengjing Hospital of China Medical University, Shenyang, China; Department of Ultrasound, Shengjing Hospital of China Medical University, Shenyang, Liaoning, China

**Author notes:** Shuangshuo Jia and Zhehan Hu have contributed equally to this work and share first authorship. **First Corresponding Author:** Yue Yang, Correspondence to: Department of Orthopedic Surgery, ShengJing Hospital, China Medical University, SanHao Street #36, HePing District, ShenYang 110000, China; Second Corresponding Author: Yang Wang, Correspondence to: Department of Ultrasound, Shengjing Hospital of China Medical University, SanHao Street #36, HePing District, ShenYang 110000, China.

**Keywords:** Osteoarthritis, cartilage intermediate layer protein (CILP), chondrocyte ferroptosis, cartilage fibrosis, exercise therapy, hyalinization of fibrocartilage

## Abstract

By analyzing the single-cell RNA-Seq libraries, the roles of cartilage intermediate layer protein (CILP) and the cartilage intermediate zone in early osteoarthritis (OA) exercise therapy were explored. An early OA rat model was established via a 4-week anterior cruciate ligament transection. The effect of moderate exercise was confirmed using histology, the open-field test, and gait analysis. The response of the cartilage intermediate zone to mechanical stimulation was explored using multiplex fluorescent immunohistochemical staining. Radiomics was used to evaluate the relatively damaged and undamaged areas in the cartilage of patients with OA. CILP was OE and KD in early OA chondrocytes, and quantitative proteomics, yeast one-hybrid, co-immunoprecipitation, Nrf2 and ubiquitination assays were used to investigate its mechanism. We found that moderate exercise upregulates CILP in the cartilage intermediate zone. CILP recovers the type II/I collagen, Sox9, and α-SMA expression ratios, and reduces Keap1-Nrf2 dimer stability, inhibiting Nrf2 ubiquitination and promoting Nrf2 nuclear translocation. Nrf2 nuclear translocation activates SLC7A11, HO-1, GPX4, and SOD-1 expression, decreases MDA content, and increases GSH content, inhibiting chondrocyte ferroptosis and promoting fibrocartilage hyalinization. In conclusion, the exercise-induced cartilage intermediate zone and CILP-Keap1-Nrf2 axis inhibit hyaline cartilage fibrosis and chondrocyte ferroptosis to alleviate early OA.

## 1. Introduction

Osteoarthritis (OA) is the most common chronic joint disease and affects approximately 530 million people worldwide.[1] Early OA may be a reversible condition,[2] and effective intervention has a profound impact on its prognosis.[3] The World Health Organization recommends exercise therapy as the first-line treatment for early OA.[4] In our previous studies, we found that moderate-intensity exercise therapy for early knee OA provided a window of opportunity to slow the OA process.[5–9] An in-depth study on exercise therapy can help develop precision medicine for treating early OA.

In our previous work, we found that cartilage and chondrocytes play important roles in maintaining the biomechanical function of the knee joint during exercise therapy.[6, 10] Articular cartilage exhibits great heterogeneity and complex microarchitecture, comprising three anatomic zones: the superficial, intermediate, and deep zones,[11, 12] which endow the tissue with biphasic mechanical properties to withstand shearing forces and compressive loading.[13] Chondrocytes in the intermediate zone were hypertrophic, slightly round in shape, and clustered together. Type II collagen fibers are randomly distributed to withstand pressure from different directions because this region is subjected to compressive and shear stresses.[14] Exercise-mediated mechanical stimuli are transmitted through the superficial zone to the intermediate zone, which plays the primary load-bearing and compression-resistance roles.[15] As a mechanosensitive zone, the cartilage intermediate zone plays a key role in early OA exercise therapy.[16]

Using single-cell transcriptome profiling to determine differential gene expression within the early OA exercise therapy, we identified a highly expressed gene that encodes cartilage intermediate layer protein (CILP), which is expressed in the intermediate zone of articular cartilage and is linked to cartilage degenerative diseases.[17, 18] In our preliminary experiments, we identified CILP as an important protein in OA treatment that affects the intermediate zone of the articular cartilage through mechanical stimulation (**Figure S1B, C**). However, the specific underlying mechanism remains unclear.

In our pseudo-temporal analysis of the pathological process in OA cartilage, cartilage fibrosis was found to play an important role in early OA (**Figure S1D, E, F, G**). Hyaline cartilage is the most common type of cartilage and has excellent mechanical properties, mainly acting as a load-bearing structure and lubricating joint movement.[19] In early OA, the articular cartilage surface is damaged, and fibrocartilage forms owing to the loss of thick collagen fibers and the formation of type I collagen.[20] The ability of fibrocartilage to bear weight and resist mechanical wear is far less than that of hyaline cartilage, and fibrocartilage in joints leads to the further progression of OA.[21] Given the anti-fibrosis effect of CILP on other diseases,[22] it is necessary to investigate the in situ transformation of fibrocartilage into hyaline cartilage via exercise-induced CILP, namely the “hyalinization of fibrocartilage” as an important principle of cartilage repair in OA.

Chondrocyte death can also lead to the excessive secretion and deposition of extracellular matrix proteins, resulting in hyaline cartilage fibrosis in early OA.[23] Ferroptosis is a programmed cell death driven by Fe^2+-^dependent lipid peroxidation.[24, 25] Our previous study showed that ferroptosis is involved in the pathogenesis and development of OA.[26] Nuclear factor erythroid 2-related factor 2 (Nrf2) is a key regulator of antioxidant responses.[27] Under oxidative stress, Nrf2 is stabilized and initiates a multistep activation pathway, including nuclear translocation.[28] The Keap1-Nrf2 pathway is critical for chondrocyte ferroptosis (**Figure S2**). It is necessary to investigate whether exercise-induced CILP affects chondrocyte ferroptosis and cartilage fibrosis via the Keap1-Nrf2 pathway.

In this study, we determined the therapeutic effects of moderate-intensity exercise on early OA. We investigated the role of the intermediate zone of articular cartilage in early OA exercise therapy. We also demonstrated the important role of exercise-induced CILP during the hyalinization of fibrocartilage. In vitro, we identified the molecular mechanism by which CILP inhibited Nrf2 ubiquitination and promoted Nrf2 nuclear translocation through the Keap1-Nrf2 signaling pathway, subsequently alleviating chondrocyte ferroptosis and promoting the hyalinization of fibrocartilage. This study highlights the potential of targeting the exercise induced CILP-Keap1-Nrf2 axis and the cartilage intermediate zone to mitigate cartilage degeneration and promote joint health.

## 2. Results

### 2.1 CILP was upregulated in cartilage of OA-affected SD rats after exercise therapy

The synovium, cartilage, and subchondral bone were analyzed using single-cell transcriptome sequencing. The t-distributed stochastic neighbor embedding (t-SNE) projection (**Figure S1B**) identified eight distinct cell clusters, including chondrocytes, fibroblasts, B cells, macrophages, and NK cells. However, this study primarily focused on the differential gene expression in chondrocytes. In the OAM group, 170 genes were downregulated, and 222 genes were upregulated compared with those in the OA group. The t-SNE projection and volcano plots (**Figure S1C**) highlighted the top 9 differentially expressed genes between the OAM and OA groups. CILP expression was upregulated in the OAM group.

### 2.2 Hyaline cartilage fibrosis is pathological change of chondrocytes in early OA

To investigate the trajectory of chondrocytes in OA further, a pseudotemporal analysis was used to simulate the pathological process in OA chondrocytes. Starting from node 1, the chondrocytes showed phenotypic changes (**Figure S1D**). We identified seven chondrocyte subtypes throughout the OA pathology, including Fc, Homc, Htc, and preFC. As shown in **Figure S1D, E**, and **F**, the Fc subtype was highly expressed in the OA group, and the cluster was in the early stage of OA chondrocytes (node 1). As shown in **Figure S1G**, we analyzed the gene expression of each chondrocyte subtype and found that the type I collagen content of the Fc subgroup increased, indicating cartilage fibrosis. Taken together, these results indicate that hyaline cartilage fibrosis is a pathological change in chondrocytes during the early stages of OA.

### 2.3 CILP is downregulated in damaged areas of articular cartilage in clinical patients

The damaged and relatively undamaged cartilaginous areas of patients with OA were divided using radiomics. Histological analyses of damaged and undamaged areas were performed in patients undergoing total knee arthroplasty (TKA). The OARSI score was used to confirm that the undamaged area had more intact cartilage than the damaged area. Using western blotting (**Figure 1B**) and immunohistochemistry (**Figure 1D**) to detect proteins in the cartilage undamaged (U) and damaged (D) regions, we found that the ratio of type II collagen to type I collagen and Sox9 expression in the U region was significantly higher than that in the D region (**Figure 1C, E).** As shown in **Figure 1F and G**, CILP was downregulated in the damaged areas of the cartilage in clinical patients.

**Figure 1.**
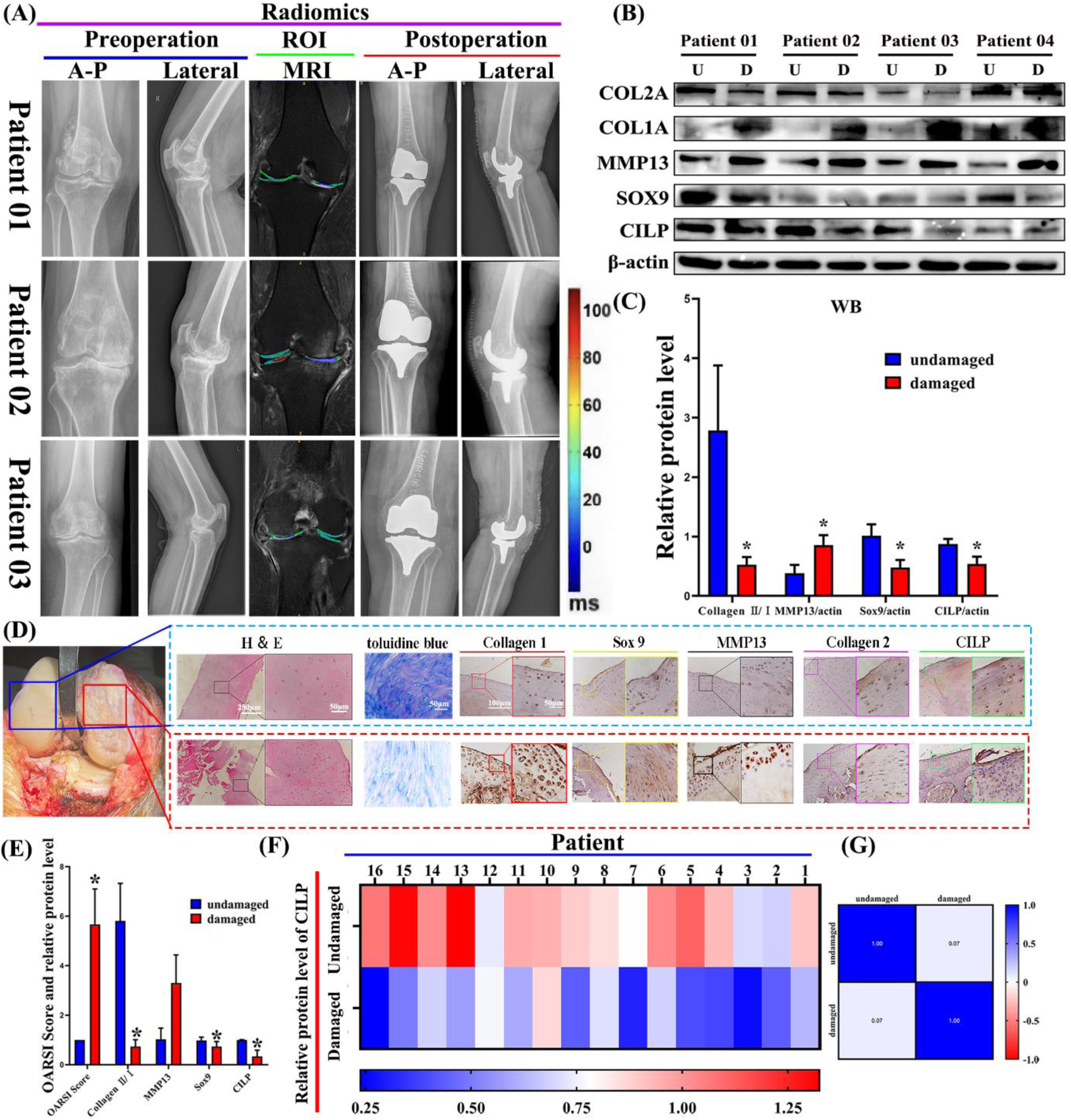
CILP is downregulated in the damaged areas of articular cartilage in clinical patients. (A) Digital radiograph and radiomics of patients with knee osteoarthritis (OA) (n = 16). Red ROI area to blue area is trend of cartilage thickness from thick to thin and from intact to damaged, anterior to posterior (A-P). (B) Western blot of CILP, typeⅡcollagen, typeⅠcollagen, Sox9, and MMP13 in undamaged (U) and damaged (D) areas. (C) Relative protein levels of CILP, type II collagen, type I collagen, Sox9, and MMP13 in undamaged and damaged areas, with β-actin as control. (D) Gross image and H&E, toluidine blue, and IHC staining of knee articular cartilage. Blue box indicates undamaged area, red box indicates undamaged area. (E) OARSI score of histology evaluation. Statistical analysis of CILP, collagen II/I, MMP13, and Sox9. \**P<* 0.001). (F) Quantification with heat maps for relative protein level of CILP in undamaged (U) and damaged (D) areas, with β-actin as endogenous control (n = 16). (G) Pearson r: correlation of CILP data.

### 2.4 Moderate-intensity treadmill exercise can increase expression of CILP in cartilage intermediate zone of SD rats and can restore imaging and histological characteristics and subjective and objective functions of knee joints in SD rats

We confirmed the occurrence of early OA in our rat model by histological evaluation. The OA group showed superficial zone edema and/or fibrillation (abrasion), focal superficial matrix condensation, or slight discontinuity (**Figure 2A**). Histological analysis based on the OARSI scores showed that the OA group of each tibia or femoral joint were between 0–12 score, which indicated early OA, and the OAM group showed therapeutic effects on the tibiofemoral joints (**Figure 2E**).

**Figure 2.**
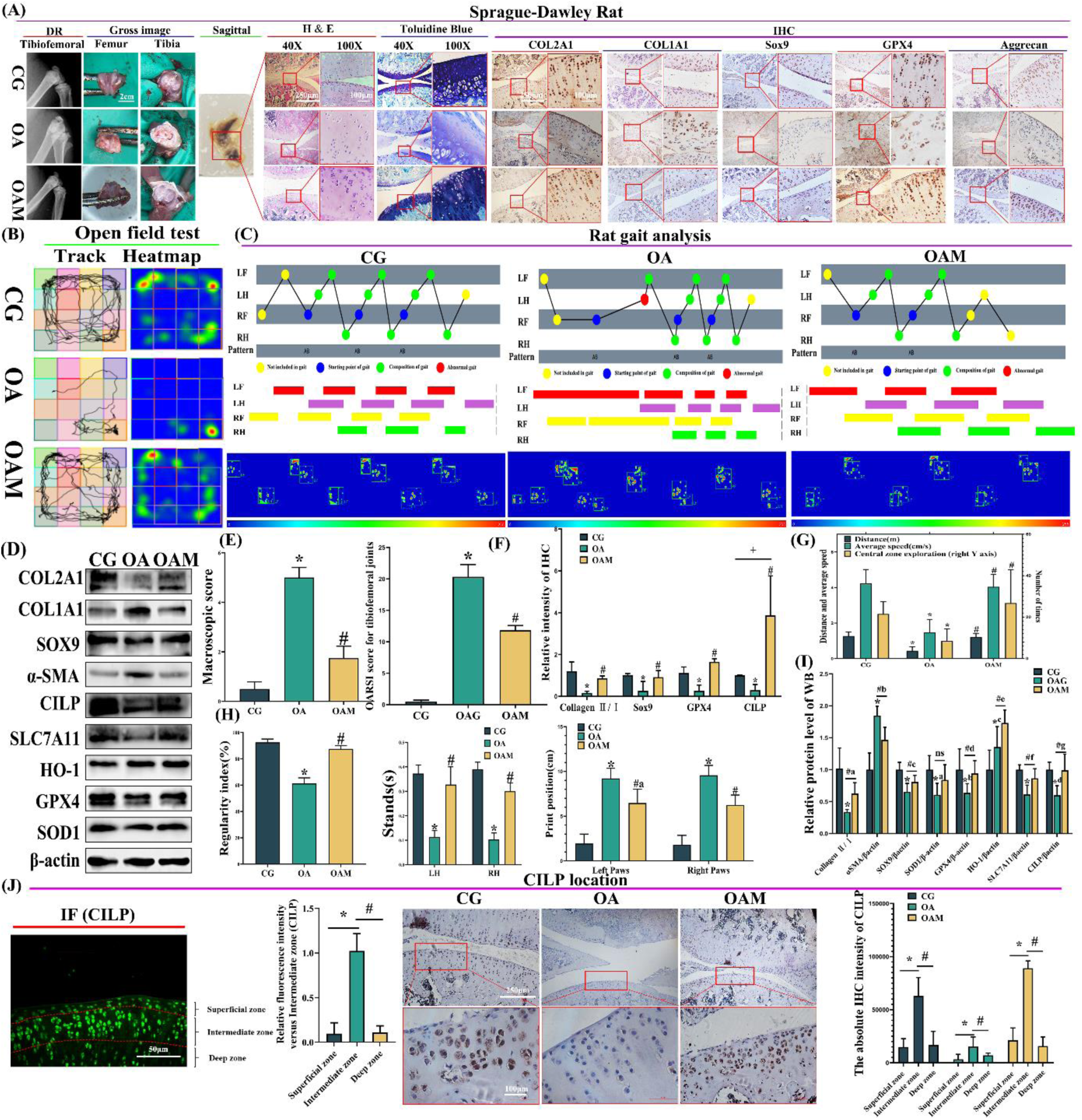
Moderate-intensity treadmill exercise can increase the expression of CILP in the cartilage of SD rats and can restore the imaging and histological characteristics and subjective and objective function of the knee joint in SD rats. (A) Digital radiograph, gross image, H&E staining, toluidine blue, and IHC evaluation of rats. (B) Track and heatmap of open field test of rats. (C) Gait analysis of rats, including step sequence, footprint time, and footprint pressure. LF: left forelimb; LH: left hindlimb; RF: right forelimb; RH: right hindlimb. (D) Western blot of rat cartilage protein expression, including CILP, typeⅡcollagen, typeⅠcollagen, Sox9,α-SMA expression, and anti-ferroptosis proteins SLC7A11, HO-1, GPX4, and SOD-1. (E) Macroscopic and OARSI scores of rats. (F) Statistical analysis of IHC evaluation of rats. (G) Open field test of rats. Left axis shows distance and average speed. Right Y axis shows central zone exploration times. (H) Statistical analysis of gait. Significant differences were found between CG and OA groups (\**P<* 0.001; ANOVA) and OA and OAM groups (#*P<* 0.001, #a P = 0.0213; ANOVA). (I) Statistical analysis of rat cartilage protein. CG and OA groups (*a P = 0.0066; *b P = 0.0178; *c P = 0.0468; *d P = 0.0022; ANOVA); OA and OAM groups (#a P = 0.0073; #b P = 0.0065; #c P = 0.0387; #d P = 0.0425; #e P = 0.0354; #f P = 0.0043; #g P = 0.0026; ANOVA). Data are expressed as the mean ± 95% confidence interval; n = 3 per group. (J) IF and IHC evaluations of CILP expression in superficial, intermediate, and deep zones in CG, OA, and OAM groups. Significant differences were found between the superficial and intermediate zone groups (\**P<* 0.001; ANOVA) and intermediate and deep zone groups (#*P<* 0.001; ANOVA).

Digital radiography revealed that the OAM group had a better articular surface and fewer osteophytes than the OA group (**Figure 2A**). Using gross images, we observed surface roughening, fibrillation, and fissures, which are typical early OA pathological changes, in the OA group compared with the CG. However, the OAM group showed alleviated symptoms compared with those of the OA group. The macroscopic score was also alleviated in the OAM group compared to that in the OA group (*P<* 0.001) (**Figure 2E**). We further evaluated the SD rats using H&E and toluidine blue staining. We found that the OA group exhibited cartilage damage and hypocellularity compared to the CG group; OAM displayed a more complete and smoother cartilage surface compared to OA. IHC staining revealed that the optical density of type II/I collagen was higher in the CG and OAM groups than in the OA group. We further noticed that Sox9, GPX4, and CILP levels were higher in patients with OAM than in those with OA.

An open-field test was used to investigate the effect of moderate-intensity exercise on the subjective recovery of knee function (spontaneous activity) in SD rats. The total distance, average speed, central zone exploration times, and motion trail were recorded (**Figure 2B**). Compared with the CG group, the OA group showed significantly reduced pain thresholds and spontaneous activity. Compared with the OA group, OAM significantly increased the total distance traveled, average speed (*P* < 0.01), and central zone exploration times (*P* < 0.01) (**Figure 2G**).

Gait analysis was performed to investigate the effect of moderate-intensity exercise on objective knee function recovery (**Figure 2C**). The regularity index and rat standing time of the left and right hind limbs were lower in the OA group than those in the CG, whereas OAM recovered these indices compared to the OA group (*P* < 0.001). The OAM group also showed a smaller paw print position than that in the OA group (*P* < 0.05). The results of the gait analysis showed that the rear limb coordination of the OAM group was greatly improved (**Figure 2H**). In the western blot, OAM recovered the ratio of type II/I collagen, Sox9, and α-SMA expression and also increased anti-ferroptosis protein SLC7A11, HO-1, GPX4, and SOD-1 expression levels (**Figure 2D, I**).

IF was used to investigate the location of CILP in the cartilage zone, and we found that the intermediate zone had the highest CILP relative fluorescence intensity (P < 0.001) (**Figure 2J**). In the IHC evaluation, the expression of CILP in the intermediate zone was highest in the CG, OA, and OAM groups (**Figure 2J**). In addition, the intermediate zone of the OAM group showed the highest CILP expression, and the OA group showed the lowest.

### 2.5 Cartilage intermediate zone acts as a mechanical-stimulation-sensitive zone in exercise therapy

Using multiplex fluorescent immunohistochemical staining (**Figure 3A, B**), we compared the differences between the zones (superficial, intermediate, and deep) and between the groups (CG, OA, and OAM) of each zone. When compared within the groups, the ratios of type II/I collagen (*P* < 0.001) and aggrecan (*P* < 0.05) were the highest in the intermediate zone. The differences in Sox9 expression were not significant (*P* > 0.05) (**Figure 3C**). As for the differences in each zone between the groups, the ratios of type II/I collagen, aggrecan, and Sox9 in the superficial, intermediate, and deep zones decreased in the OA group but recovered in the OAM group (*P* < 0.05) (**Figure 3D, E, F**).

**Figure 3.**
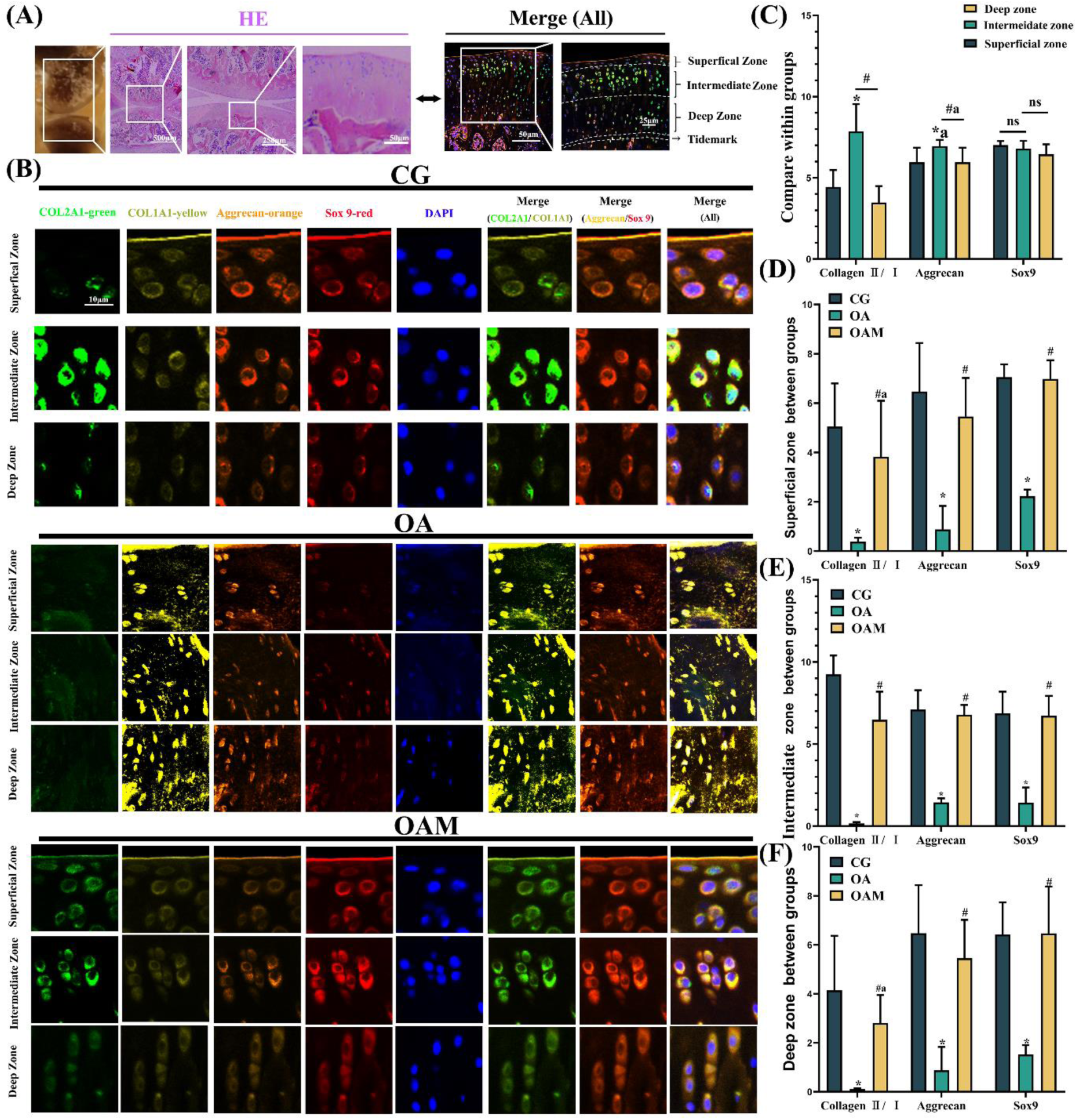
The cartilage intermediate zone acts as a mechanical-stimulation-sensitive zone in exercise therapy. (A) Using multiplex fluorescent immunohistochemical staining, we divided the cartilage into three zones (superficial, intermediate, and deep). (B) Multiple fluorescent immunohistochemical staining of CG, OA, and OAM groups; green: collagen II; yellow: collagen I; aggrecan: orange; red: sox9; blue: DAPI. (C) Statistical analysis of the relative fluorescence intensity differences between zones (superficial, intermediate, and deep). Significant differences were found between superficial and intermediate zones (\**P<* 0.001; *a P = 0.033; ANOVA) and intermediate and deep zones (#*P<* 0.001, #a P = 0.045; ANOVA). (D) Statistical analysis of relative fluorescence intensity of superficial zone between groups. Significant difference comparison gave the same result as in Figure 2. #a P = 0.0041. (E) Statistical analysis of relative fluorescence intensities of intermediate zones between groups. (F) Statistical analysis of relative fluorescence intensities of deep zones between groups. Significant differences were found between the CG and OA groups (\**P<* 0.001; ANOVA) and OA and OAM groups (#*P<* 0.001, #a P = 0.0014; ANOVA). Data are expressed as mean ± 95% confidence interval; n = 3 per group.

### 2.6 Generation 3 chondrocytes (G3) could mimic early fibrosis of articular cartilage with early OA

During chondrocyte passage, the adherent morphology of the chondrocytes gradually changed from completely irregular polygons to slender spindle shapes. P3 cells showed the early appearance of fibrochondrocytes (**Figure 4A**). The type II/I collagen, aggrecan, and Sox9 ratio decreased during passaging (**Figure 4B**). Compared to primary chondrocytes, the type II/type I collagen ratio decreased from 5.85 to 1.63 in Generation 3 chondrocytes (G3) (**Figure 4D**). As shown in **Figure 4C and E**, the relative fluorescence intensity of type II collagen in G3 was significantly lower than that in G0 (*P* < 0.001). By comparing the cell modeling using the inducer TGF-β1 for cartilage fibrosis, we found that the type II/I collagen, Sox9, and α-SMA levels of G3 cells were not significantly different from those in the early cartilage fibrosis model induced by low concentrations of TGF-β1 (10 ng/mL) (**Figure 4F, H**).

**Figure 4.**
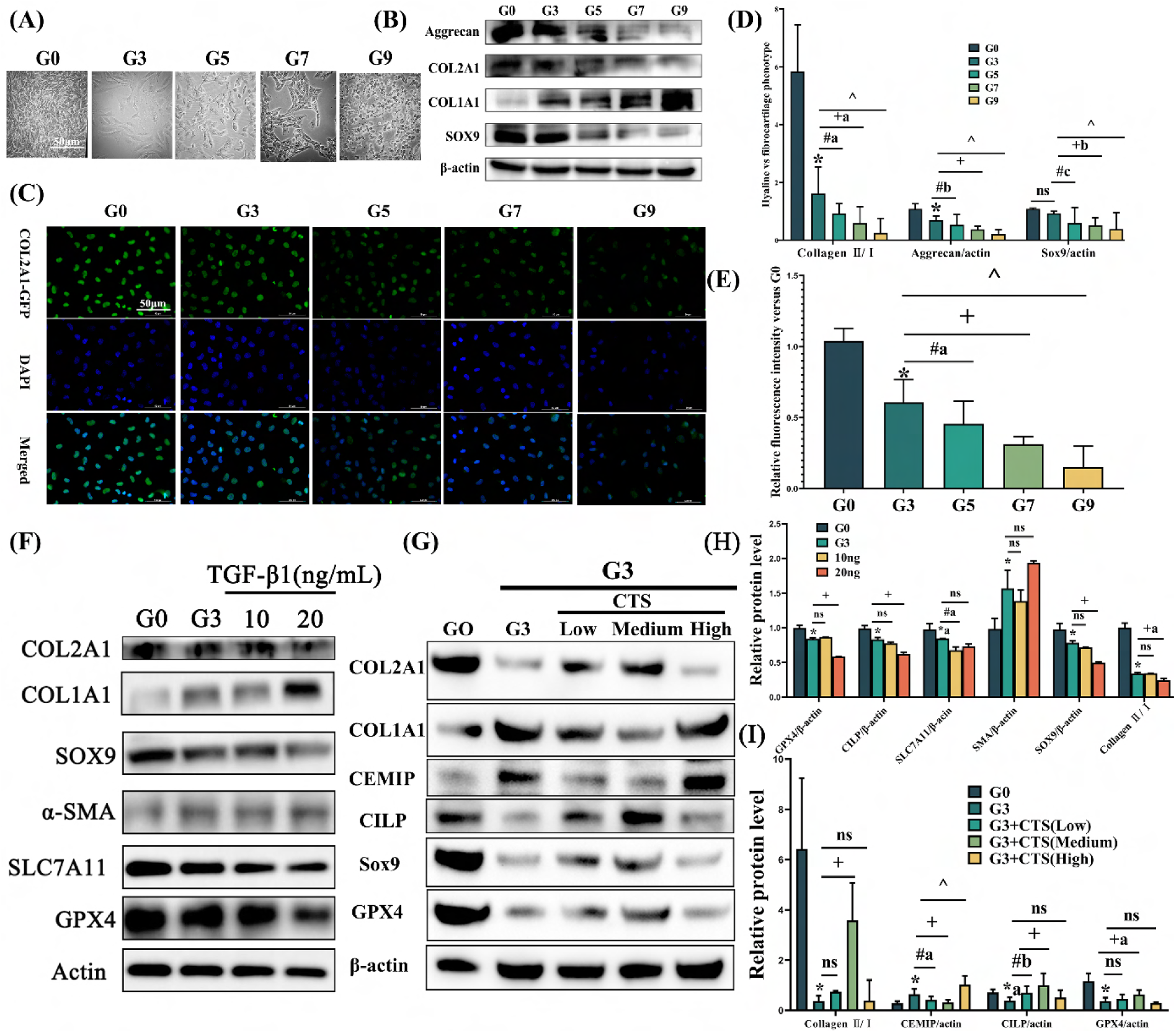
Generation 3 chondrocytes (G3) could mimic early fibrosis of articular cartilage with early OA, and moderate CTS alleviates early fibrosis of chondrocytes by upregulating CILP. (A) Gross image of adherent morphology of chondrocytes during passage. (B) Western blot of hyaline cartilage versus fibrocartilage phenotype in different passage chondrocytes. G0: primary chondrocyte; G3: Generation 3 chondrocyte; and so on. (C) Immunofluorescence of collagen II in different passage chondrocytes. (D) Statistical analysis of the hyaline cartilage versus fibrocartilage phenotype in different passage chondrocytes. Significant differences were found between G0 and G3 (\**P<* 0.001; ANOVA), G3 and G5 (#*P<* 0.001, #a P = 0.019; #b P = 0.021; #c P = 0.004; ANOVA), G3 and G7 (+*P<* 0.001; +a P = 0.002; +b P = 0.003; ANOVA), and G3 and G9 groups (^*P<* 0.001; ANOVA). (E) Statistical analysis of relative fluorescence intensity in different passage chondrocytes (#a P = 0.015; ANOVA). (F) Western blot of expression of hyaline cartilage versus fibrocartilage phenotype in G3 chondrocytes and TGF-β1-induced chondrocyte fibrosis models. (G) Western blot analysis of phenotypic proteins such as typeⅡcollagen, typeⅠcollagen, CEMIP, CILP, Sox9, and GPX4 of G3 chondrocytes after CTS treatment. (H) Statistical analysis of G3 chondrocytes and TGF-β1-induced chondrocyte fibrosis models. G0 and G3 groups (\**P<* 0.001; *a P = 0.0052; ANOVA); G3 and TGF-β1 10 ng groups (#*P<* 0.001; #a P = 0.0122; ANOVA); G3 and TGF-β1 20 ng group (+*P<* 0.001; +a P = 0.0293; ANOVA). (I) Statistical analysis of G3 chondrocytes after CTS treatment. G0 and G3 groups (\**P<* 0.001; *a P = 0.006; ANOVA); G3 and G3+CTS (low) groups (#*P<* 0.001; #a P = 0.007; #a P = 0.008; ANOVA); G3 and CTS (medium) groups (+*P<* 0.001; +a P = 0.0015; ANOVA); G3 and CTS (high) groups (^*P<* 0.001; ANOVA). Data are expressed as mean ± 95% confidence interval; n = 3 per group.

### 2.7 Moderate CTS alleviates early fibrosis of chondrocytes in vitro by upregulating CILP

The upregulation of CILP was most significant in G3 cells after moderate CTS treatment (*P* < 0.001). Moreover, the protein expression levels of type II/I collagen, Sox9, and GPX4 showed a significant recovery compared to those of the G3 early fibrosis phenotype (*P* < 0.05). The fibrosis characteristic protein CEMIP was higher in G3, especially in the G3+high-CTS group (*P* < 0.001).

### 2.8 CILP competitively binds to Keap1 protein sites and reduces stability of Keap1-Nrf2 dimer, decreases Nrf2 ubiquitination, and promotes Nrf2 nuclear translocation

To investigate the molecular mechanisms underlying the effects of CILP on chondrocytes, we performed a proteomic analysis of intracellular protein expression following CILP overexpression. Volcano plots showed the differentially expressed proteins (**Figure S2A**), selected according to the criteria of (log2 |fold-change| ≥ 1.2 and *P<* 0.05). The Keap1 protein was enriched in the OE-CILP group compared to the G3 group (**Figure S2B**). Subcellular localization showed that Keap1 was enriched in the cytoplasm of chondrocytes (**Figure S2C**). Following quantitative proteomics, we utilized GO functional annotation and KEGG pathway enrichment analyses to explore the potential signaling pathways interacting with Keap1. Based on the protein function cluster (**Figure S2D**), we found that CILP interacted with Keap1 and affected Nrf2, affecting ferroptosis and promoting early OA cartilage fibrosis (**Figure S2E, F**).

In our study, the potential binding sites for KEAP1 and CILP were identified using yeast one-hybrid assays. CILP promotes KEAP1/NRF2 dissociation by competing with Nrf2 for binding to the Arg483 site of KEAP1. Co-IP results showed a more positive interaction between CILP and Keap1 in the OE-CILP group than in the KD-CILP group. As shown in **Figure 5B**, the Keap1 protein was detected in IP-CILP in the OE-CILP group, and no Keap1 protein was observed in the band of the IgG-binding protein. In addition, IB-CILPs were detected in the IP-Keap1 group (**Figure 2C**). Co-immunoprecipitation results also showed that the interaction between Keap1 and Nrf2 was decreased in the OE-CILP group. As shown in **Figure 5D**, Nrf2 protein was detected in the IP-Keap1 in the KD-CILP group, and less Nrf2 protein was observed in the band of the OE-CILP group binding protein (**Figure 2D**). Furthermore, the trend of IB-Keap1 expression detected in the IP-Nrf2 group was similar to that described above (**Figure 2E**). As shown in **Figure 5F**, Nrf2 ubiquitination decreased in the OE-CILP group, while Nrf2 ubiquitination increased in the KD group. In addition, OE-CILP accumulated NRF2 that had translocated to the nucleus. However, KD-CILP reversed this trend (**Figure 5G**).

**Figure 5.**
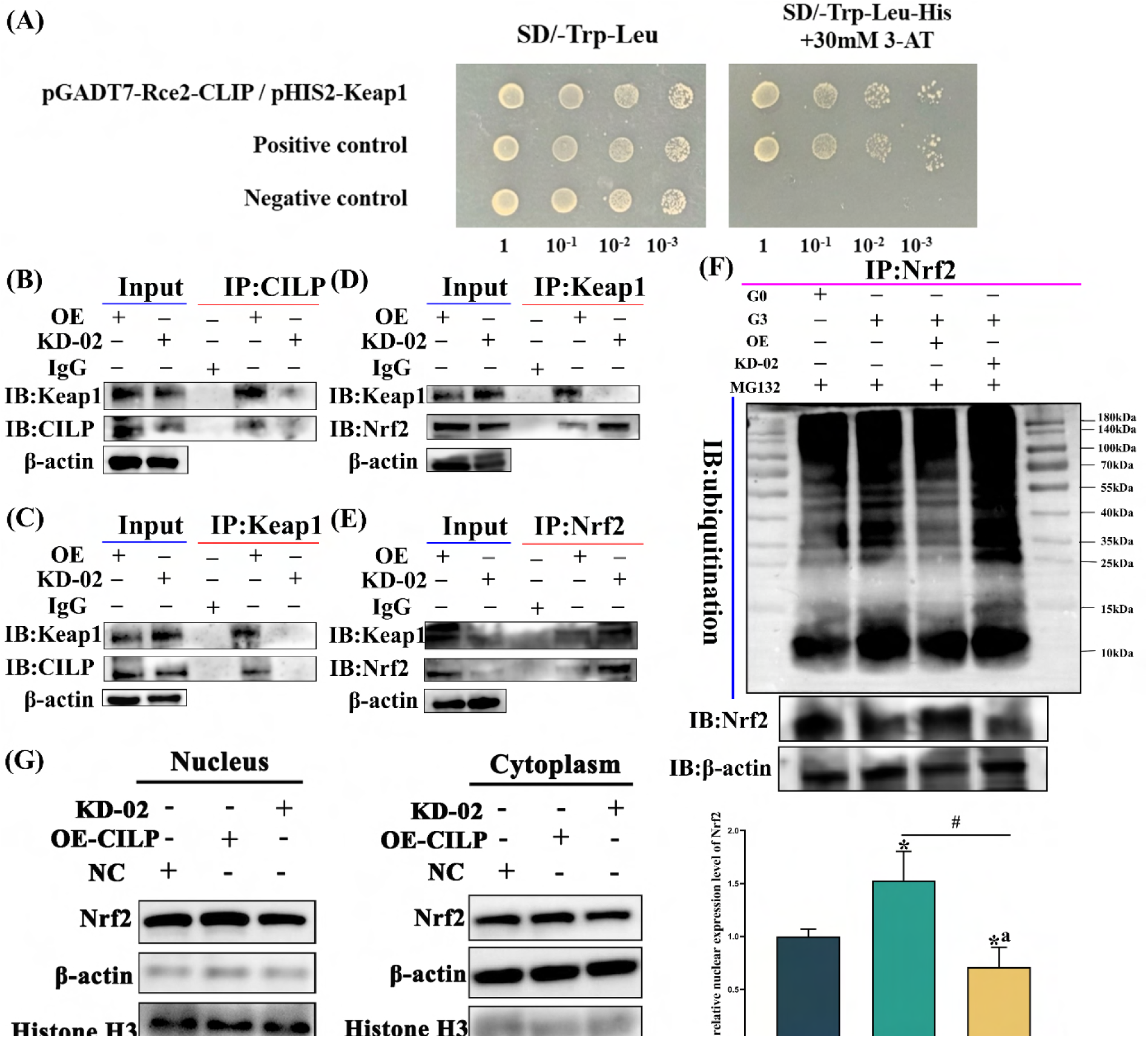
CILP competitively binds to sites of Keap1 protein and reduces stability of Keap1-Nrf2 dimer, decreasing Nrf2 ubiquitination and promoting Nrf2 nuclear translocation. (A) Yeast one-hybrid assays to detect potential binding sites for Keap1 and CILP. (B) OE/KD-CILP group CILP immunoprecipitated from chondrocytes with anti-Keap1 and anti-CILP antibodies, respectively, was analyzed using immunoblotting. (C) OE/KD-CILP group Keap1 immunoprecipitated from chondrocytes with anti-Keap1 and anti-CILP antibodies, respectively, were analyzed using immunoblotting. (D) OE/KD-CILP group Keap1 immunoprecipitated from chondrocytes with anti-Keap1 and anti-Nrf2 antibodies, respectively, were analyzed using immunoblotting. (E) OE/KD-CILP group Nrf2 immunoprecipitated from chondrocytes with anti-Keap1 and anti-CILP antibodies, respectively, were analyzed using immunoblotting. (F) OE/KD-CILP group Nrf2 immunoprecipitated from chondrocytes with anti-ubiquitination antibodies. (G) Western blot of Nrf2 in nucleus and cytoplasm of chondrocytes in OE/KD-group. Significant differences were found between negative control and OE-CILP, KD-02 (\**P<* 0.001; *a P = 0.0099; ANOVA), and OE-CILP and KD-02 groups (#*P<* 0.001; ANOVA). β-actin and Histone H3 were used for negative control.

### 2.9 CILP ameliorates chondrocyte ferroptosis through Keap1-Nrf2 pathway in vitro

Using western blotting, we found that the OE-CILP group increased the ratio of type II/I collagen, Sox9, and α-SMA expression and also increased the expression levels of anti-ferroptosis proteins SLC7A11, HO-1, GPX4, and SOD-1 (**Figure 6A, B**). However, these therapeutic effects could be reversed by KD-01, especially KD-01 (**Figure 6C, D**). Confocal microscopy revealed a decrease in intracellular ROS levels in the OE-CILP group. Flow cytometric analysis supported this result (**Figure 6E**). There was also an increased mitochondrial membrane potential (**Figure 6G**) and decreased intracellular Fe2+ levels in the OE-CILP group (**Figure 6I**). In addition, TEM showed ferroptosis in the CILP-knockdown group, mainly characterized by smaller mitochondria, increased membrane density, and reduced mitochondrial cristae. OE-CILP ameliorated chondrocyte ferroptosis (**Figure 6J**). The levels of MDA (**Figure 7A**) and GSH (**Figure 7B**) in chondrocytes were also significantly decreased via CILP (*P* < 0.05). These results suggest that CILP overexpression reverses the progression of cartilage fibrosis and chondrocyte ferroptosis in early OA. KD-CILP reversed these therapeutic effects.

**Figure 6.**
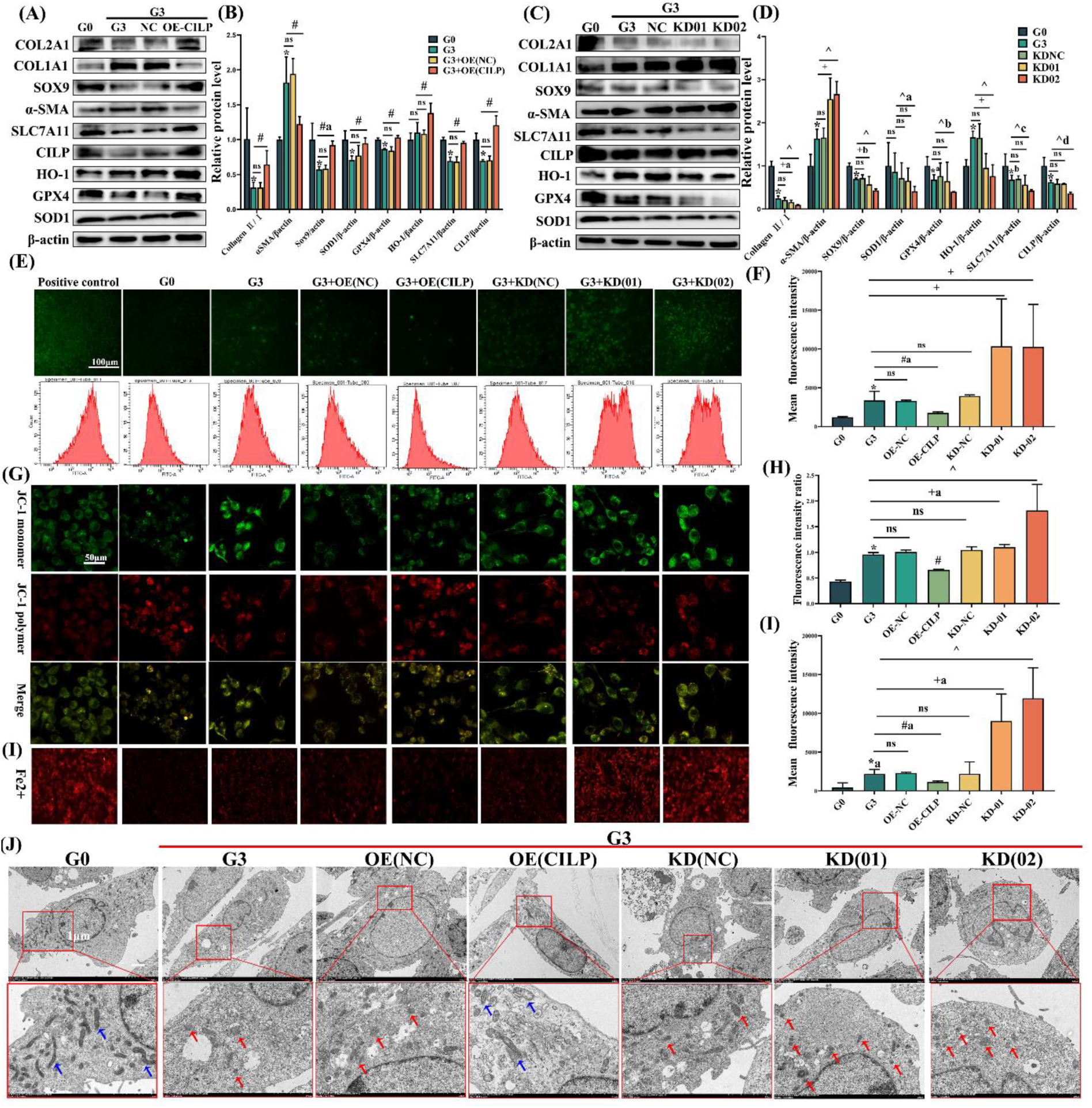
CILP ameliorates chondrocyte ferroptosis through Keap1-Nrf2 pathway in vitro. (A) Western blot of OE –CILP G3 chondrocyte, including CILP*P<P<*, typeⅡcollagen, typeⅠcollagen, Sox9, α-SMA expression, and anti-ferroptosis protein SLC7A11, HO-1, GPX4, and SOD-1 expression. (B) Statistical analysis of proteins of G3 chondrocytes after OE-CILP treatment in G0 and G3 groups (\**P<* 0.001; ANOVA); G3 and OE-CILP groups (#*P<* 0.001; #a P = 0.041; ANOVA). (C) Western blot of KD–CILP (KD-01, KD-02) G3 chondrocyte protein expression, including CILP, typeⅡcollagen, typeⅠcollagen, Sox9, α-SMA expression, and anti-ferroptosis proteins SLC7A11, HO-1, GPX4 and SOD-1. (D) Statistical analysis of G3 chondrocytes after KD-01, KD-02 treatment. G0 and G3 (\**P<* 0.001; *a P = 0.0271; *b P = 0.0013; ANOVA); G3 and KD-01 (+*P<* 0.001; +a P = 0.002; +b P = 0.0146; ANOVA); and G3 and KD-02 groups (^*P<* 0.001; ^a P = 0.0313; ^b P = 0.0498; ^c P = 0.0068; ^d P = 0.0015; ANOVA). (F) Confocal microscope showed intracellular ROS levels, and flow cytometry analysis showed cellular ROS result. (F) Statistical analysis of G3 chondrocytes after OE-CILP, KD-01, and KD-02 treatment. G0 and G3 (\**P<* 0.001; ANOVA); G3 and OE-CILP (#*P<* 0.001; #a P = 0.011; ANOVA); G3 and KD-01, and KD-02 groups (+*P<* 0.001; ANOVA). (G) Results of incubation with JC-1 fluorescent and FerroOrange probes (I) and corresponding statistical analysis. (H) Statistical analysis of relative fluorescence intensity of JC-1 (+a P = 0.041; ANOVA). (I) Statistical analysis of relative fluorescence intensity of Fe^2+^ (*a P = 0.008; #a P = 0.0015; +a P = 0.0012; ANOVA). Data are expressed as mean ± 95% confidence interval; n = 3 per group. (J) Transmission electron microscopy of mitochondrial morphology changes in chondrocytes. Blue arrows indicate normal mitochondria, red arrows indicate ferroptosis-changed mitochondria (a reduction in mitochondrial volume, disappearance of cristae, and membrane disruption).

**Figure 7.**
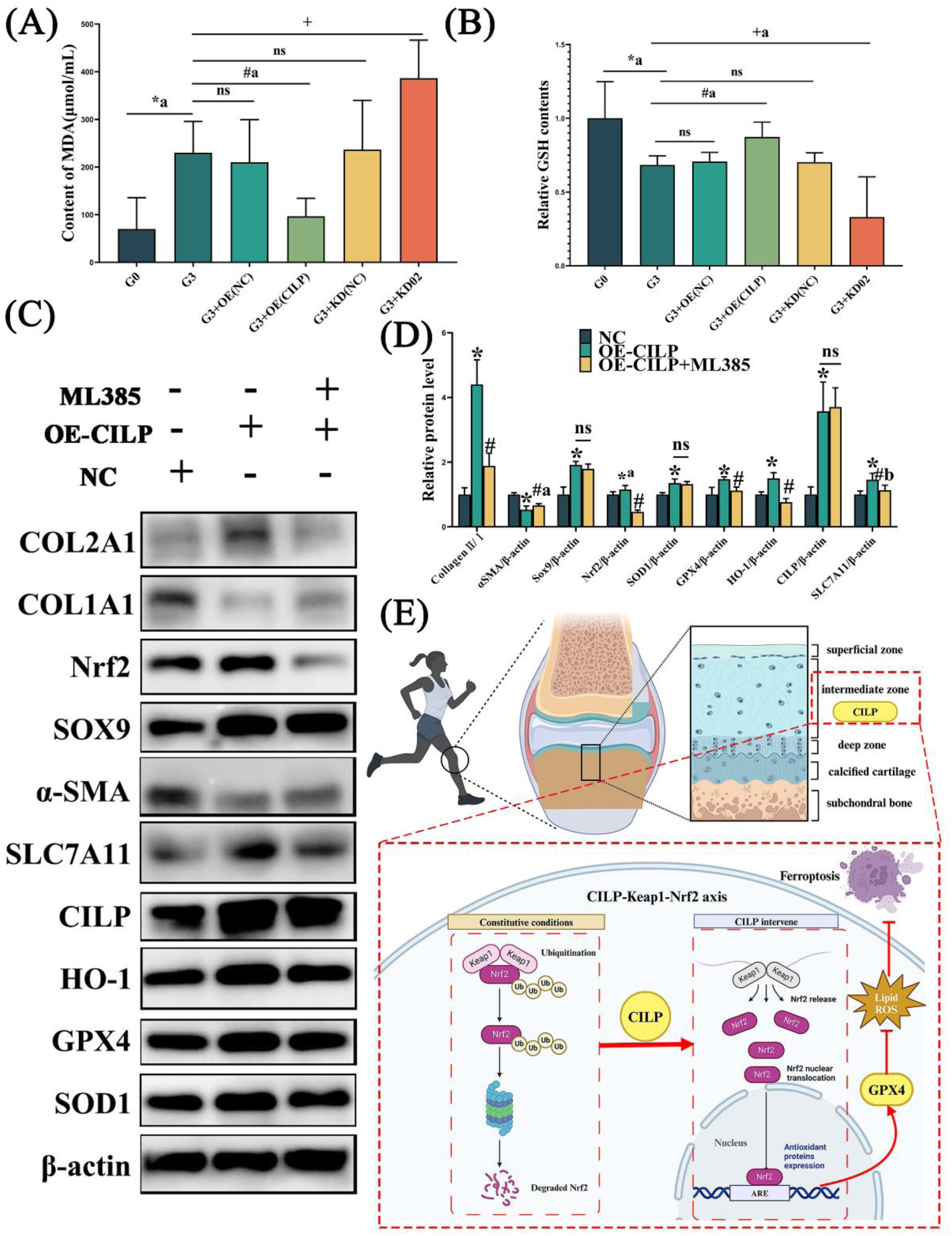
Exercise-CILP-Keap1-Nrf2 axis ameliorates early OA cartilage fibrosis by inhibiting chondrocyte ferroptosis. (A) Content of Malondialdehyde (MDA) in chondrocytes. Significant differences were found between G0 and G3 (\**P<* 0.001; *a P = 0.0018; ANOVA); OE-CILP and G3 (#*P<* 0.001; #a P = 0.0016; ANOVA); and G3 and KD-02 groups (+*P<* 0.001; ANOVA). (B) Content of GSH in chondrocytes. Significant differences were found between G0 and G3 (\**P<* 0.001; *a P = 0.006; ANOVA); OE-CILP and G3 (#*P<* 0.001; #a P = 0.0023; ANOVA); and G3 and KD-02 groups (+*P<* 0.001; +a P = 0.0056; ANOVA). (C) Western blot of ML385 as ferroptosis activator to antagonize anti-fibrotic and anti-ferroptosis effects of CILP. (D) Statistical analysis of relative protein expression after ML385 treatment. Significant differences were found between NC and OE-CILP (\**P<* 0.001; *a P = 0.0018; ANOVA) and OE-CILP and OE-CILP +ML385 groups (#*P<* 0.001; #a P = 0.0264; #a P = 0.042; ANOVA). (E) Exercise-induced cartilage intermediate zone and CILP-KEAP1-Nrf2 axis inhibit hyaline cartilage fibrosis and chondrocyte ferroptosis to alleviate early OA.

### 2.10 Exercise-CILP-Keap1-Nrf2 axis ameliorates early OA cartilage fibrosis by inhibiting chondrocyte ferroptosis

This study investigated the importance of the exercise-CILP-Keap1-Nrf2 axis and ferroptosis in treating early OA cartilage fibrosis. We used ML385, an Nrf2 inhibitor, to antagonize the anti-fibrotic and anti-ferroptotic effects of CILP. The ratio of type II/I collagen (*P* < 0.001) and α-SMA (*P* < 0.05) were significantly decreased by ML385 interference. There were no statistically significant differences in the Sox9, CILP, or SOD-1 levels. The levels of anti-ferroptosis proteins, such as Nrf2, GPX4, HO-1, and SLC7A11, decreased significantly (*P* < 0.05).

## 3. Discussion

This study confirmed that moderate-intensity exercise has a therapeutic effect on early OA in terms of histology, animal behavior (open field test), and subjective and objective knee function (gait analysis), highlighting the positive role of the articular cartilage intermediate zone in response to mechanical stimulation in exercise therapy for early OA. We are the first to report that moderate-intensity exercise upregulates CILP in the articular cartilage intermediate zone to promote hyalinization of the fibrocartilage and alleviate early OA. We also observed that CILP competitively bound to Keap1 protein and reduced the stability of the Keap1-Nrf2 dimer, reducing the degree of Nrf2 ubiquitination and promoting Nrf2 nuclear translocation. Nrf2 nuclear translocation activates SLC7A11, HO-1, GPX4, and SOD-1 expression, decreases MDA content, and increases GSH content in chondrocytes, ultimately inhibiting chondrocyte ferroptosis. Finally, we conclude that the exercise-induced CILP-Keap1-Nrf2 axis mitigates early OA through fibrocartilage hyalinization and suppressing chondrocyte ferroptosis.

This study focused on early OA. The early symptoms of OA are relatively insidious but gradually worsen with disease progression.[29] Reduced patient compliance owing to pain leads to limited treatment strategies. Early OA is a reversible process that is less affected by pain, and personalized exercise prescriptions can be developed.[2] To establish the early OA model, we followed Pritzker et al. and Maier et al.,[30, 31]who identified early OA with the OARSI score (0–12 each tibia or femoral side), matrix staining depletion within the upper third of the cartilage, and reduction of the cartilage thickness and collagen content. The 4-week ACLT was used to establish an early OA rat model, which followed the above criteria and showed a decrease in cartilage type II collagen and loss of cartilage aggrecan.[32–34] Chondrocytes are vulnerable to dedifferentiation during in vitro passage, leading to fibrosis and the mechanical degradation of newly formed cartilage.[35] We used passaged chondrocytes that were dedifferentiated through cultivation, which was evident from the loss of proteoglycans, decreased COL-II deposition, and increased synthesis of COL-I compared to primary chondrocytes (G0).[36–38] Chondrocytes shift from round to fibroblastic shape with increasing passage number. Several molecular differentiation markers have been identified. Among them, type II/type I collagen, aggrecan, and Sox9 levels decrease during the dedifferentiation process and are often used to assess chondrogenic loss (**Figure 4**).

Compared with primary chondrocytes, the ratio of type II/type I collagen decreased from 5.85 to 1.63 in Passage 3 chondrocytes (G3). By comparing the cell modeling using the inducer TGF-β1 [39–41] for joint fibrosis, we found that P3 cells were close to the mild cartilage fibrosis model induced by low concentration TGF-β1. The use of dedifferentiation induced by the passage of chondrocytes to mimic the fibrosis of early OA cartilage is also an innovative aspect of this study.

In our pseudo-temporal analysis of the pathological process in OA cartilage, disorders in the synthesis and catabolism of chondrocytes led to cartilage damage and degeneration. Hyaline chondrocytes begin to proliferate actively, transform, or dedifferentiate into cells with a fibroblast phenotype and form cell clusters in the cartilage lesion area to adapt to changes in the cartilage microenvironment. Fibrotic cartilage is characterized by dramatic changes in cell shape, metabolism, and cytoskeletal structure, resulting in the increased synthesis of type I collagen (collagen I), which impairs the biological function of cartilage tissue by forming fibrocartilage without mechanical capacity.[20] Decreased expression of type II collagen and increased expression of type I collagen are the main features of articular chondrocyte dedifferentiation and fibrosis.[19, 21] Moreover, chondrocyte death and the abnormal catabolism and metabolism of the cartilage matrix caused by ferroptosis can lead to cartilage defects and damage. During cartilage repair after injury, the appearance of fibrotic cartilage can further aggravate the symptoms of OA. In this study, we used a ferroptosis activator to interfere with the therapeutic effect of CILP and found that ferroptosis activation aggravated cartilage fibrosis (**Figure 7C**). Combined with previous studies,[23, 42] the joint study of chondrocyte ferroptosis and cartilage fibrosis after cartilage injury was the focus of the present study.

In this study, the cartilage was found to be heterogeneous and anisotropic, with three regions having different compositions and structural mechanical properties. Chondrocytes in the superficial zone were more likely to exhibit a fibrocartilage phenotype due to the higher amount of frictional type I collagen caused by frequent joint activity (**Figure 3)**. Chondrocytes in the superficial area were oblate and scattered on the cartilage surface. Type II collagen fibers are laterally distributed to withstand wear and stress (**Figure 3B**).[43] Lubrication and painless sliding of the joint occur in the superficial zone, whereas the intermediate zone transfers the load applied in the superficial zone to the deep zone of the cartilage.[12] Collagen fibers in the deep zone are arranged perpendicularly to the articular surface (**Figure 3B**).[43, 44] The intermediate zone accounts for approximately 40–60% of articular cartilage thickness; chondrocytes are rounded and medium in shape, and collagen fibers are arranged randomly and perpendicular to the articular surface. The main matrix components in the intermediate zone are type II collagen (Col2a1) and aggrecan (**Figure 3B**).[13] Moreover, the aggrecan content in the MZ reached the maximum level, and collagen type II was highly expressed and randomly distributed to withstand pressures from different directions (**Figure 3B**), as this region was subjected to compressive and shear stresses.

We established a moderate-intensity treadmill exercise protocol for exercise therapy for early OA, according to our previous studies.[5, 6, 8, 45] We used histological analysis, an open field test, and gait analysis to confirm the therapeutic effect of moderate exercise on early OA in terms of physiological, psychological, subjective, and objective knee function recovery (**Figure 2**). Single-cell transcriptome sequencing was performed on the entire knee joint after exercise, and it was found that exercise may affect early OA through CILP in the intermediate zone of the cartilage (**Figure S1**). In the CILP localization experiment (**Figure 2J**), we confirmed that CILP was mainly enriched in the intermediate zone and was upregulated by exercise. In the intermediate zone, the ratio of type II/I collagen to aggrecan also significantly increased (*P* < 0.05). As a mechanical stress-sensitive zone, the intermediate zone receives mechanical stimulation signals from exercise therapy.

Using proteomic and bioinformatic analyses, we found that CILP affected hyaline cartilage fibrosis and chondrocyte ferroptosis via Keap1. In our study, the potential binding of Keap1 to CILP was identified using yeast one-hybrid assays (**Figure 5A**). CILP promotes Keap1/Nrf2 dissociation by competing with Nrf2 to bind to Keap1, decreasing Nrf2 ubiquitination and accumulating Nrf2 translocation to the nucleus. Our current study indicated that the CILP-Keap1-Nrf2 antioxidative signaling pathway is involved in fibrocartilage hyalinization. Hyaline cartilage phenotypes, such as the ratios of type II collagen to type I collagen, Sox9, and α-SMA, were upregulated after CILP intervention. Oxidative stress and the antioxidant system are critical for fibrosis.[46, 47] We also demonstrated that Nrf2-mediated anti-ferroptotic activity depends on the induction of SLC7A11, GPX4, HO-1, and SOD1, which are currently recognized as central repressors of ferroptosis.[48, 49] This alleviated ferroptosis in chondrocytes by reducing the accumulation of ROS and inhibiting iron overload and lipid peroxidation in the mitochondria (**Figure 6**). Taken together, these results indicated that the CILP-Keap1-Nrf2 axis alleviated OA by inhibiting chondrocyte ferroptosis and cartilage fibrosis.

Our study has some limitations. First, the intermediate zone of cartilage as a mechanosensitive zone has not been studied in depth, and we did not extract cartilage in the intermediate zone separately from animal and patient cartilage for analysis. Second, histological and radiographic evaluations of early OA have not yet been clearly established. We used the OARSI scoring system and histological evaluation to eliminate bias as much as possible. Third, the association between hyaline cartilage fibrosis and ferroptosis in chondrocytes has only been preliminarily confirmed and will be further investigated in sequential studies.

## 4. Conclusion

In conclusion, exercise alleviates ferroptosis in chondrocytes and promotes the hyalinization of fibrocartilage by mechanically stimulating the intermediate zone of the cartilage and regulating the CILP-Keap1-Nrf2 axis in this region. This study highlights the different roles of different cartilage subzones in the treatment of early OA using exercise and clarifies the mechanism by which CILP in the mechanosensitive zone (intermediate zone) hyalinizes fibrocartilage in early OA through intracellular signal transduction, which may provide a theoretical basis and reference for exercise therapy for OA.

## Declaration

### Author contributions

Conceptualization: SJ, YY, ZH; Methodology: SJ, ZH, CN, YS, GY, ZZ; Formal analysis and investigation: ZL, WZ, ZZ; Writing - original draft preparation: SJ, ZH; Writing - review and editing: SJ, YY, YW; Funding acquisition: YY; Resources: YW, YY; Supervision: YY

## Data availability statement

Data will be made available on request.

## Funding statement

This study was supported by the National Natural Science Foundation of China (Grant No. 82102613), the Liaoning Provincial Science and Technology Plan Project (Doctoral Research Initiation Project) (Grant No. 2023012135-JH3/4500) and the Technology Talent Project of Liaoning Provincial Education Department (Grant No. LJKQZ2021028).

## Conflict of interest disclosure

The authors have no conflicts of interest to declare that are relevant to the content of this article.

## Ethics approval statement

The studies involving human participants were reviewed and approved by the protocol and experiments on joint specimen after human replacement were approved by the Ethics Committee of Shengjing Hospital of China Medical University (No: 2021PS265K). The animal study was reviewed and approved by the Ethics Committee of China Medical University (no. 2021PS220K).

## Patient consent statement

Written informed consent was obtained from the individual(s) for the publication of any potentially identifiable images or data included in this article.

## 5. Experimental Section/Methods

### 5.1 Participants

The protocol and experiments on joint specimen after human total knee arthroplasty were approved by the Ethics Committee of Shengjing Hospital of China Medical University (No: 2019PS629K) which abides the principles set out in the World Medical Association Helsinki Declaration. Inclusion/exclusion criteria of study participants had been described in previous literature. [5, 7, 10] We obtained informed consent from all of our patients.

### 5.2 Radiomics

An Avanto 3.0T MR Scanner (SIEMENS MAGNETON), was used to collect patients’ image with an 8-channel coil for the knee joint. Radiomics was used to predict and classify the cartilage damaged and undamaged areas in MRI images of OA patients. Following the protocol of the previous study, [50, 51] we briefly stated the steps below. All pretreatment MRI images were acquired. Two experienced radiologists (7-year experience in bone and joint MRI reading) manually delineated the region of interest (ROI) using ITK-SNAP (version 3.8.0; http://www.itksnap.org/). The ROI cover the edge of articular cartilage on coronal T1WI and coronal fat-suppressed PDWI images to generate a three-dimensional ROI of articular cartilage. The feature extraction was performed using the Standardized Environment for Radiomics Analysis (SERA). [50] All the features were normalized using Z-score normalization. The intraclass correlation coefficient (ICC) was used to evaluate the consistency of the extracted image features. Extractions of 623 features showed good reproducibilities (ICCs >0.80). Using maximum relevance minimum redundancy (mRMR) and least absolute shrinkage and selection operator (LASSO) to reduce the dimensionality of the texture features in MRI. The rad-score was calculated according to the feature weight.

Then, two support vector machine (SVM) models were built using the selected radiomics features to classify the damaged and undamaged regions. Different kernel functions of SVM were tested to find the best performance kernel functions, including linear, radial basis function (RBF), cubic and sigmoid kernel functions. A five-fold cross-validation method was used to train and validate the model.

### 5.3 Human OA articular cartilage and clinicopathological features collection

As mentioned in our previous study, [5, 10] the cartilage and the clinicopathological features of patients with knee osteoarthritis after total knee replacement were collected and divided into damaged and undamaged groups according to radiomics result and the Osteoarthritis Research Society International (OARSI) scale, which uses a point system ranging from 0 to 24. [52] We compared the damaged area with the undamaged area in the same donors.

### 5.4 Experimental Animals and Early Osteoarthritis Model

Male Sprague–Dawley (SD) rats (140 ± 5 g, 4-week-old, specific-pathogen-free, eight rats per group, and six rats per cage) were procured from HFK Bioscience Co. Ltd. (Beijing, China). The study was approved by the Ethics Committee of China Medical University (no. 2021PS130K(X1)), adhering to the regulations stipulated in the World Medical Association Helsinki Declaration. The upkeep and care of the experimental rats complied with the committee’s guidelines, as described in previous studies. [5, 8]

The OA model was established using Anterior Cruciate Ligament Transection (ACLT). [53] The early OA model was established over a period of 4 weeks, following previous studies and confirmed by histology evaluation. [53] The 36 rats were segregated into three groups: the control group (CG), the osteoarthritis group (OA), and the osteoarthritis plus moderate exercise group (OAM), with 12 rats in each group. Post-modelling, the OAM group began treadmill training on the ZH-PT animal treadmill exercise platform (Zhongshidichuang Science & Technology Development Co. Ltd., Beijing, China) using appropriate light, acoustic, and electrical stimuli. The moderate treadmill exercise protocol (19.2 m/min) was based on our prior studies. [5, 8, 54, 55] As our previous research demonstrated that moderate-intensity exercise has the optimal therapeutic effect on osteoarthritis. [5, 6, 8, 9, 54, 56] A total of 36 SD rats were used for single-cell transcriptome sequencing (n=12).

### 5.5 Tissue Sampling and Collection

After the final treadmill exercise session, rats were euthanized via a pentobarbital overdose, as per our established protocol. [5, 8] Tissue samples, including synovium, cartilage, and subchondral bone, were subsequently collected for single-cell transcriptome sequencing. For histological and immunohistochemical evaluations, knee joints were fixed using a 4% paraformaldehyde solution (Sigma-Aldrich, St. Louis, MO, United States) for a duration of 7 days. Subsequently, the samples underwent decalcification in a 20% EDTA solution (Sigma-Aldrich, St. Louis, MO, USA) for 6 weeks. Following decalcification, the samples were dehydrated using a graded series of ethanol and xylene (Sigma-Aldrich) and embedded in paraffin (Sigma-Aldrich). [5]

### 5.6 Single-cell Transcriptome Sequencing and Bioinformatics Analysis

Synovium, cartilage, and subchondral bone were collected from the CG, OA, and OAM groups. Sequencing libraries were generated following a comprehensive transcriptome sequencing and single-cell gene-expression profiling protocol as previously reported. [57, 58] Single-cell RNA-Seq libraries were generated using the SeekOne® Digital Droplet Single Cell 3’ library preparation kit (SeekGene, Beijing, China).

To identify exercise-related differentially expressed genes in chondrocyte, we compared the expression profiles of the chondrocyte of the OA, and OAM groups. Volcano plots of differentially expressed genes were generated using the Limma/R package (version 3.5.1). The primary parameters were set as log fold change, with a log fold change >2 indicating differentially expressed genes. These differentially expressed genes were then submitted to the DAVID v6.8 tool (https://david.ncifcrf.gov/) for annotation, visualization, and integrated discovery. [5]

### 5.7 Pseudo-time trajectory

Following by the previous study, [59] the Monocle2 R package (v2.14.0) (http://cole-trapnell-lab.github.io/monocle-release/) was used to dissect gene expression changes associated with cell state transitions in chondrocyte clusters. The ‘DDRTree’ function in Monocle2 R package was used to reduce the dimensionality of the datasets, the ‘orderCells’ function was used to sorted cells in a pseudo-temporal order. Pseudo-time trajectories with default parameters were inferred using the ‘reduceDimension’ function, and the reconstructed trajectories were visualized using the ‘plot_cell_trajectory’ function. Significantly different marker genes were extracted from each module, where clustering was performed based on pseudo-temporal expression patterns and representative heat maps were generated.

### 5.8 Histology evaluation of articular cartilage of human and early OA animal models

We embedded cartilage of human and joints from SD rats in paraffin. Sagittal sections of 4.5-μm thickness were cut from the tibiofemoral joints and stained with hematoxylin and eosin (H&E), toluidine blue, for histological evaluation as described in our previous study. [5, 6] As detailed in our earlier research, [5, 10] we categorized the cartilage and the clinicopathological characteristics of patients with knee osteoarthritis into damaged and undamaged area. This categorization was guided OARSI, which uses a point system ranging from 0 to 24. [52]

In animal models, knee joint images were captured by X-ray (MX-20, Faxitron X-ray, Corp., Lincolnshire, IL, United States) and NRecon version 1.6 software (Bruker). As detailed in our previous study, both the tibial and femoral joints were evaluated based on macroscopic score and OARSI score. [5, 60] Both the tibial and femoral joints were evaluated based on a maximum OARSI score of 48. Followed by the previous study, [31] we use score 0-12 of 24 (tibial or femoral) to identify early OA.

### 5.9 Immunohistochemistry

Briefly, following the kit manufacturer’s instructions (CAT # SP-9001, Zhongshan Jinqiao, China), we performed immunostaining of proteins using a two-step method. Procedures were carried out as we have previously described. [9, 10] Antibodies including, anti-Collagen II (ab34712, 1:200, abcam, Cambridge, MA, USA), anti-Collagen I (ab270993, 1:200, abcam), anti-Sox9 (ab185966, 1:200, abcam), anti-MMP13 (ab219620, 1:200, abcam), anti-CLIP (sc-12725, 1:200, Santa Cruz Biotechnology, Santa Cruz, CA, USA), anti-GPX4 (ab125066, 1:200, abcam).

### 5.10 Open field test

The open field apparatus was obtained from Shanghai Xinruan Software Technology (China). The experimental animals were quickly and gently placed in the central area of the experimental box (100 cm × 100 cm × 60 cm black open-field box), and the video acquisition and analysis system was started at the same time to automatically record the activity of the experimental animals in the open field. The experimental time was usually set to 6 min. At the end of each test, the feces of the previous animal should be removed, and the odor should be removed with 75% alcohol, and the chamber should be dry and odorless before the next animal experiment.The data were exported for statistical analysis. The distance (cm) the rat moved within the cage during 6 min, average speed, and number of explorations in the central region was calculated.

### 5.11 Rat gait analysis

The CatWalk gait analysis system (Noldus Information Technology B.V., Netherlands) was used to evaluate the gait dynamics of SD rats. On the day after the end of the exercise treatment, the CatWalk system was used to test rats in each group. Gait analysis software (CatWalk XT 10.6) was used to calculate gait parameters automatically. This experiment mainly evaluated the effects of the following gait parameters after OA exercise treatment, including the regularity index (%), posture (stands) and hind limb fingerprint position (cm).

### 5.12 Multiple fluorescent immunohistochemical staining

After paraffin sections were deparaffinized, the slides were washed with sterilized water for 1 min and repeated three times. A five-color multiplex immunofluorescence histochemical kit (absin, abs50013, China) was used to evaluate SD rat joint section. Following the manufacturer’s instructions, enzymatic antigen retrieval (C1033, Solarbio Science & Technology Co., Ltd., Beijing, China) was performed at 37°C for 30 min to repair antigens in the sections. Then, 3% H_2_O_2_ was used at 25°C for 30 min to eliminate endogenous peroxidase activity, while blocking serum (15019, Cell Signaling Technology, Danvers, MA, United States) was used at 25°C for 30 min to block nonspecific antigens. Then, sections were incubated with anti-Collagen II (ab34712, 1:50, abcam), anti-Collagen I (ab270993, 1:5, abcam), anti-Sox9 (ab185966, 1:50, abcam), anti-Aggrecan (abs121273, 1:50,absin).

Fluorescence staining was used to amplify the signal (different colors) for the color display of the corresponding antibody. After each round of staining, the staining was confirmed by fluorescence microscope, then cleaned with TBST, and new primary antibodies were added. Nuclei were stained with 4,6-diamidino-2-phenylindole (DAPI) for 5 min. Coverslips were visualized under a confocal microscope (Olympus).

### 5.13 Primary Chondrocyte Isolation, Culture, Passage and Treatment

Rat primary chondrocytes were isolated from the knee articular cartilage of 4-week-old SD rats. The 3 mg/ml pronase K (V900887, Sigma-Aldrich, St. Louis, MO, United States) and 2.0 mg/ml collagenase D (C0130, Sigma-Aldrich) were used to cartilage digestion. The isolation and culture of chondrocyte were detailed in our previous study. [5, 8] Resuspended chondrocytes were transferred into culture flasks and cultured with DMEM/F-12 supplemented with 10% FBS (abs972, Absin Bioscience Inc., Shanghai, China) and 1% penicillin with streptomycin (Hyclone Laboratories Inc., Logan, UT, United States) at 37°C in a 5% CO2 incubator. Then, the chondrocytes were passaged at a 1:4 ratio with 0.05% trypsin (C0202; Beyotime Biotech, Shanghai, China) until 70%–80% confluence. The primary(1), 3, 5, 7 and 9 passages chondrocytes were collected for characteristic phenotype (cell morphology, Collagen II/ I) analysis, and the most suitable early OA chondrocyte fibrosis model was established followed by previous study. [61]

The role of CILP in OA treatment was assessed by transfecting G3 chondrocytes with recombinant adenovirus carrying either CILP knockdown or overexpression sequences (multiplicity of infection [MOI] = 30, polybrene = 1 μg/mL; Hanbio, China). CILP knockdown conditions were grouped as follows: sh-NC (KD-NC), HBAD-Adeasy-r-CILP shRNA1-EGFP (KD-01), and HBAD-Adeasy-r-CILP shRNA2-EGFP (KD-02). CILP overexpression conditions were divided into Ad-NC (OE-NC) and HBAD-Adeasy-r-CILP - 3xflag-EGFP (OE-CILP). In rescue experiment, ML385 (HY-100523, MedChem Expres, NJ, USA) was used as ferroptosis acativator.

### 5.14 Exposure of chondrocytes to cyclic tensile strain (CTS)

Chondrocytes were grown on collagen I-coated Bioflex 6-well culture plates (Flexcell International, Hillsborough, NC) to70%-80% confluence. CTS experiments were performed using the FX-5000 Flexcell system (Flexcell International, McKeesport, PA). As detailed by our previous study, [7, 9] low (5%, 0.5 Hz), moderate (10%, 0.5 Hz) and high (15%, 0.5 Hz) CTS were applied to chondrocyte.

### 5.15 Western Blot Analysis

As detailed in our previous study, [5, 6, 8, 10] both cartilage and chondrocytes were lysed using RIPA (9806S, Cell Signaling Technology). The antibodies including: anti-Collagen II (ab34712, 1:1500, abcam), anti-Collagen I (ab270993, 1:1500, abcam), anti-Sox9 (ab185966, 1:1500, abcam), anti-α-SMA (SAB5500002, 1:1500, Sigma-Aldrich, St.Louis, MO,USA), anti-SLC7A11(ab307601, 1:1500, abcam), anti-CEMIP (21129-1-AP, 1:1500, proteintech, Wuhan, China), anti-GPX4(ab125066, 1:1500, abcam), anti-Aggrecan (13880-1-AP, 1:1500, proteintech), anti-CILP (sc-12725, 1:1500, Santa Cruz Biotechnology), anti-HO-1(ab189491, 1:1500, abcam), anti-SOD1(10269-1-AP, 1:1500, proteintech), anti-Nrf2 (ab62352, 1:1500, abcam), anti-β-actin(ab8227, 1:1500, abcam).

### 5.16 Quantitative Proteomics of CILP-treated chondrocyte and Related Bioinformatics Analysis

To examine the protein profiles of the CILP-treated chondorcyte, we employed the data-independent quantitative proteomics analysis. [62] All analyses were conducted using a Q-Exactive HFX mass spectrometer (Thermo, USA) equipped with a Nanospray Flex source (Thermo, USA). Chromatographic separation was carried out on the EASY-nLC 1000 HPLC System (Thermo, USA). The parameters were followed by previous study, [63] all of the Q Exactive raw data was searched using DIA-NN (v1.8.1). A global false discovery rate (FDR) was set to 0.01, and protein groups were considered for quantification if they had at least 2 peptides. Finally, the differences between each group were analyzed using bioinformatics analysis.

### 5.17 Yeast One-Hybrid Assay

The Keap1 enhancer was used as bait to screen for its binding factors from a cDNA library derived from chondrocytes using the BD Matchmaker One-Hybrid System (Clontech, K1617-1) and the manufacturer suggested protocol with modifications. The coding sequences of *CILP* and *Keap1* were inserted into the XhoI and BamHI restriction sites of the pGADT7-Rec2 vector (Supplementary Data Table S1).

The constructs were co-transformed into yeast strain Y187. Yeast colonies were grown at 30°C for 3-5 days on SD/−Trp/−Leu medium (DDO). The stock solution and bacterial solution diluted 10, 100 and 1000 times (1, 10^-1^, 10^-2^,10^-3^) were then screened on SD/−Trp/−Leu/−His medium (TDO) supplemented with 30 mM 3-amino-1,2,4-triazole (3-AT) for 3 days. p53HIS2 and pGADT7-p53 were used as positive controls. p53HIS2 and pGADT7-SmMYBs were used as the corresponding negative controls.

### 5.18 Co-Immunoprecipitation (Co-IP)

For Co-Immunoprecipitation (Co-IP), cells were lysed using IP/Co-IP lysis buffer (PC102, Epizyme, Shanghai, China). The lysates were subsequently immunoprecipitated using anti-KEAP1 antibody (4678, 1:50, Cell Signaling Technology, Danvers, MA, USA), anti-Nrf2 (ab62352, 1:50, abcam), anti-CILP (sc-12725, 1:50, Santa Cruz Biotechnology). A total of 2μg rat IgG antibody (A7031, Beyotime, Shanghai, China) was used as an internal control. These components were mixed with Protein A/G magnetic beads (HY-K0202, MedChemExpress, Monmouth Junction, NJ, USA) and incubated at 4℃ with rotation overnight. The following steps were the same as those used in western blot analysis.

### 5.19 Nrf2 ubiquitination assay

The chondrocyte lysates were immunoprecipitated using anti-Nrf2 (ab62352, 1:50, abcam), and then incubated with ubiquitylation assay kit (ab139467, abcam) at 37°C for 1 – 4 hours followed by manufacturer’s instructions. Then analyze by SDS-PAGE and Western blot with anti-ubiquitination antibody

### 5.20 Cytosol-nuclei fractionation

We used the nuclear and cytoplasmic protein extraction kit to dissociate the cytoplasmic and nuclear proteins (PK10014, Proteintech, China) according to the manufacturer’s instructions. Fractions were analyzed by SDS-PAGE and western blot with specific antibodies.

### 5.21 Cellular ROS Production

Chondrocytes were seeded in 6-well plates (1.5×10^6^ cells/well). ROS production was measured using 2′, 7′-dichlorodihydrofluorescein diacetate (DCFH-DA) (S0033, Beyotime), which is directly oxidized by ROS (e.g., superoxide ion, hydrogen peroxide, and hydroxyl). Chondrocytes were incubated with 10μM DCFH-DA for 35 min at 37°C in the dark. Fluorescence was detected by fluorescent microscopy and measured with the BD FACSCalibur (488 nm, excitation; 525 nm, emission).

### 5.22 JC-1 staining

The mitochondrial membrane potential in chondrocyte was determined via a JC-1 fluorescent probe (abs50016, absin), and incubated with JC-1 working solution for 20 min at 37 °C. Then, JC-1 buffer solution was used to wash the cells at least three times. The fluorometric ratio of JC-1 aggregates to JC-1 monomers was used as an indicator of mitochondrial dysfunction.

### 5.23 Ferrous ion detection

The level of cellular iron was detected using FerroOrange (Dojindo, Kumamoto, Japan) following the manufacturer’s instructions and imaged using a fluorescence microscope. We measured the fluorescence intensity using ImageJ.

### 5.24 Malondialdehyde (MDA) and GSH detection

MDA content and GSH content were detected using the Malondialdehyde assay kit (abx257171, Abbexa Ltd, Cambridge, UK), reduced glutathione assay kit (abx150350, Abbexa Ltd) following the manufacturer’s instructions. The level of GSH was measured using a microplate reader at the absorption wavelengths of 405 nm. The values for the levels of MDA was measured using a spectrophotometer at the absorption wavelengths of 532 nm.

### 5.25 Transmission electron microscopy (TEM)

Following by our previous study, [5, 7, 8] the cellular morphology and subcellular structures were then observed using a Hitachi 800 transmission electron microscope (TEM) (Tokyo, Japan).

### 5.26 Statistical Analysis

Results are presented as means ± 95% confidence interval for difference (mean± 95% CI), analyzed using GraphPad Prism 5 (GraphPad Software Inc, San Diego, California, USA). Statistical evaluations were conducted using Student’s t-test and one-way ANOVA, facilitated by IBM SPSS Statistics 25.0. A p-value of < 0.05 was considered statistical significant.

## Supplementary material

**Supplementary Figure 1.**
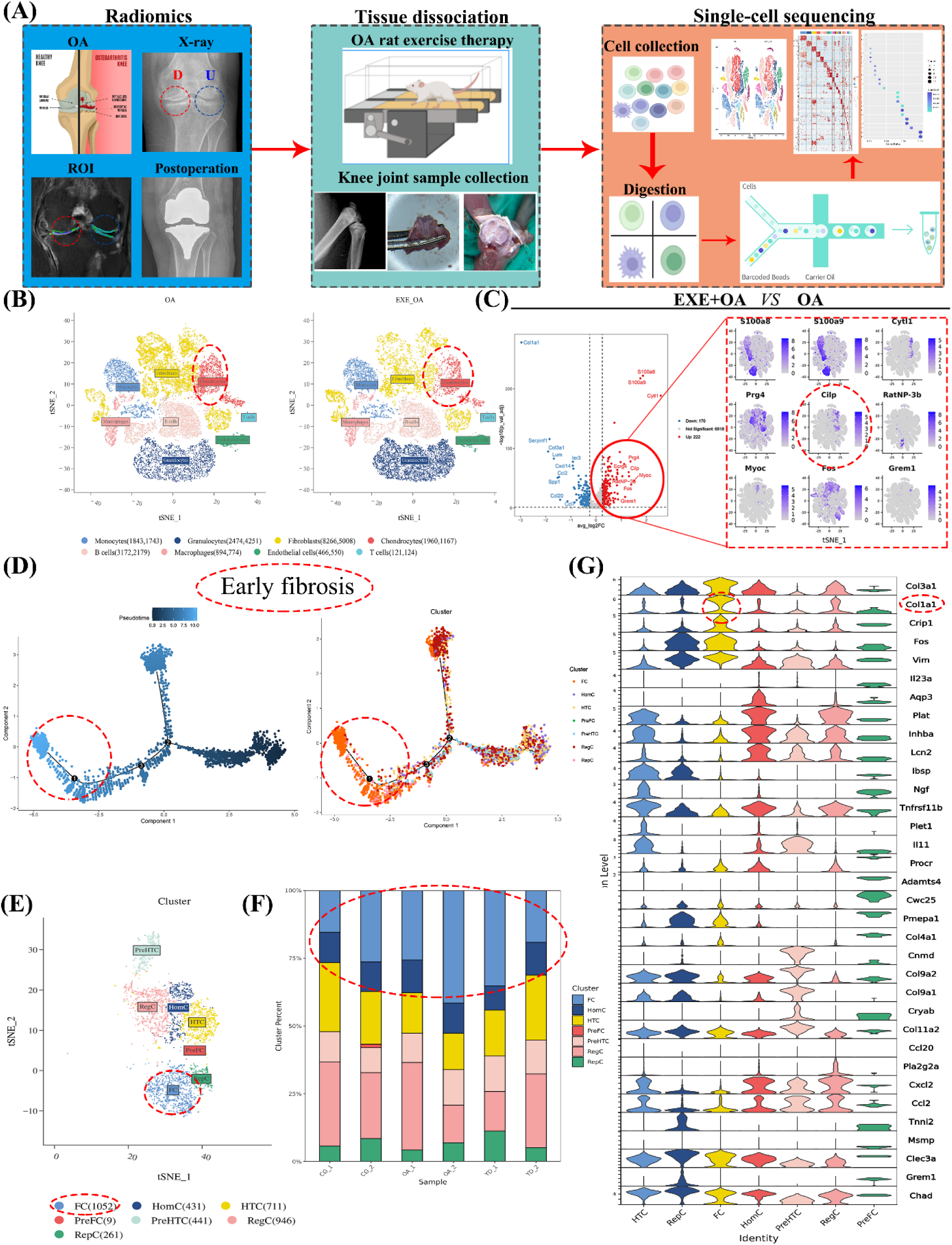
CILP and hyaline cartilage fibrosis are crucial in chondrocytes during early OA, with radiomics, single-cell transcriptomic, pseudo-time trajectory, and chondrocyte proteomic analysis. (A) Radiomics, single-cell transcriptomics, and chondrocytes proteomics diagram. (B) t-distributed stochastic neighbor embedding (t-SNE) of cell type of whole joint of SD rats between OAM and OA groups. (C) Differentially expressed genes in chondrocytes between OAM and OA groups were displayed in volcano plot and t-SNE. (D) Pseudo-temporal analysis of pathological process of OA chondrocytes. Starting from node 1, chondrocytes showed some phenotypic changes. (E) Identified chondrocyte subtypes throughout OA pathology; (F) Fc subtype was more expressed in OA group, and cluster was in early stage of OA chondrocytes. (G) Gene expression of each subtype of chondrocytes and type Ⅰ collagen content of Fc subgroup were increased, indicating cartilage fibrosis.

**Supplementary Figure 2.**
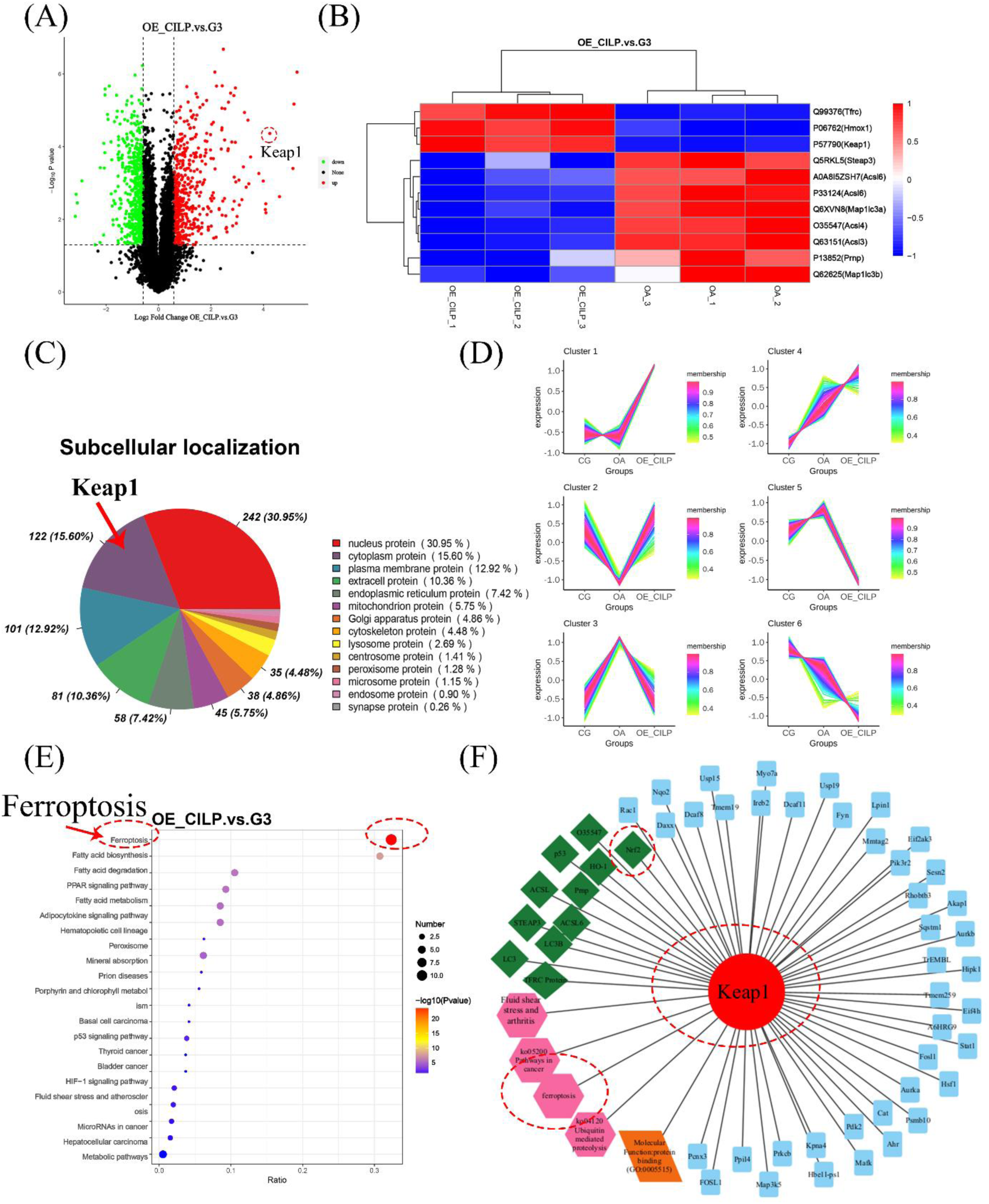
Proteomic analysis of CILP in treatment of early OA through Keap1-Nrf2 and chondrocyte ferroptosis. (A) Differentially expressed protein in chondrocytes between OE-CILP and G3 groups displayed in volcano plot. (B) Bioinformatic analysis of differential proteins associated with ferroptosis (displayed in heatmap). (C) Subcellular location showed Keap1 enriched in chondrocyte cytoplasm. (D) Differentially expressed protein cluster analysis. (E) Bioinformatic analysis of enrichment function of different proteins between OE-CILP and G3 groups. (F) Differentially enriched protein Keap1 interacts with string network of ferroptosis.

## Notes

### Competing Interest Statement

The authors have declared no competing interest.

## References

[1] M. Englund, Osteoarthritis, part of life or a curable disease? A bird’s-eye view, J Intern Med 293(6) (2023) 681–693 10.1111/joim.13634

[2] R. Paz-González, V. Balboa-Barreiro, L. Lourido, V. Calamia, P. Fernandez-Puente, N. Oreiro, C. Ruiz-Romero, F.J. Blanco, Prognostic model to predict the incidence of radiographic knee osteoarthritis, Annals of the rheumatic diseases 83(5) (2024) 661–668 10.1136/ard-2023-225090

[3] G.A. Hawker, L.S. Lohmander, What an earlier recognition of osteoarthritis can do for OA prevention, Osteoarthritis Cartilage 29(12) (2021) 1632–1634 10.1016/j.joca.2021.08.007

[4] T. Moseng, T.P.M. Vliet Vlieland, S. Battista, D. Beckwée, V. Boyadzhieva, P.G. Conaghan, D. Costa, M. Doherty, A.G. Finney, T. Georgiev, M. Gobbo, N. Kennedy, I. Kjeken, F.P.B. Kroon, L.S. Lohmander, H. Lund, C.D. Mallen, K. Pavelka, I.A. Pitsillidou, M.P. Rayman, A.T. Tveter, J.E. Vriezekolk, D. Wiek, G. Zanoli, N. Østerås, EULAR recommendations for the non-pharmacological core management of hip and knee osteoarthritis: 2023 update, Ann Rheum Dis (2024) 10.1136/ard-2023-225041

[5] S. Jia, Y. Yang, Y. Bai, Y. Wei, H. Zhang, Y. Tian, J. Liu, L. Bai, Mechanical Stimulation Protects Against Chondrocyte Pyroptosis Through Irisin-Induced Suppression of PI3K/Akt/NF-κB Signal Pathway in Osteoarthritis, Frontiers in cell and developmental biology 10 (2022) 797855 10.3389/fcell.2022.797855

[6] Z. Li, Z. Huang, H. Zhang, J. Lu, Y. Tian, S. Piao, Z. Lin, L. Bai, Moderate-intensity exercise alleviates pyroptosis by promoting autophagy in osteoarthritis via the P2X7/AMPK/mTOR axis, Cell Death Discov 7(1) (2021) 346 10.1038/s41420-021-00746-z

[7] P. Shen, S. Jia, Y. Wang, X. Zhou, D. Zhang, Z. Jin, Z. Wang, D. Liu, L. Bai, Y. Yang, Mechanical stress protects against chondrocyte pyroptosis through lipoxin A4 via synovial macrophage M2 subtype polarization in an osteoarthritis model, Biomed Pharmacother 153 (2022) 113361 10.1016/j.biopha.2022.113361

[8] J. Liu, S. Jia, Y. Yang, L. Piao, Z. Wang, Z. Jin, L. Bai, Exercise induced meteorin-like protects chondrocytes against inflammation and pyroptosis in osteoarthritis by inhibiting PI3K/Akt/NF-kappaB and NLRP3/caspase-1/GSDMD signaling, Biomed Pharmacother 158 (2022) 114118 10.1016/j.biopha.2022.114118

[9] Y. Yang, Y. Wang, Y. Kong, X. Zhang, H. Zhang, Y. Gang, L. Bai, Mechanical stress protects against osteoarthritis via regulation of the AMPK/NF-kappaB signaling pathway, J Cell Physiol 234(6) (2019) 9156–9167 10.1002/jcp.27592

[10] Z. Lin, J. Miao, T. Zhang, M. He, Z. Wang, X. Feng, L. Bai, JUNB-FBXO21-ERK axis promotes cartilage degeneration in osteoarthritis by inhibiting autophagy, Aging Cell 20(2) (2021) 10.1111/acel.13306

[11] Y. Wu, B. Ayan, K.K. Moncal, Y. Kang, A. Dhawan, S.V. Koduru, D.J. Ravnic, F. Kamal, I.T. Ozbolat, Hybrid Bioprinting of Zonally Stratified Human Articular Cartilage Using Scaffold-Free Tissue Strands as Building Blocks, Adv Healthc Mater 9(22) (2020) e2001657 10.1002/adhm.202001657

[12] D. Dehghan-Baniani, B. Mehrjou, P.K. Chu, W.Y.W. Lee, H. Wu, Recent Advances in “Functional Engineering of Articular Cartilage Zones by Polymeric Biomaterials Mediated with Physical, Mechanical, and Biological/Chemical Cues”, Adv Healthc Mater 12(10) (2023) e2202581 10.1002/adhm.202202581

[13] C.A. Tee, J. Han, J.H.P. Hui, E.H. Lee, Z. Yang, Perspective in Achieving Stratified Articular Cartilage Repair Using Zonal Chondrocytes, Tissue Eng Part B Rev 29(3) (2023) 310–330 10.1089/ten.TEB.2022.0142

[14] Y. Jia, H. Le, X. Wang, J. Zhang, Y. Liu, J. Ding, C. Zheng, F. Chang, Double-edged role of mechanical stimuli and underlying mechanisms in cartilage tissue engineering, Front Bioeng Biotechnol 11 (2023) 1271762 10.3389/fbioe.2023.1271762

[15] L. Yin, Y. Wu, Z. Yang, V. Denslin, X. Ren, C.A. Tee, Z. Lai, C.T. Lim, J. Han, E.H. Lee, Characterization and application of size-sorted zonal chondrocytes for articular cartilage regeneration, Biomaterials 165 (2018) 66–78 10.1016/j.biomaterials.2018.02.050

[16] T.J. Mosher, Y. Liu, C.M. Torok, Functional cartilage MRI T2 mapping: evaluating the effect of age and training on knee cartilage response to running, Osteoarthritis Cartilage 18(3) (2010) 358–64 10.1016/j.joca.2009.11.011

[17] Z. Yao, H. Nakamura, K. Masuko-Hongo, M. Suzuki-Kurokawa, K. Nishioka, T. Kato, Characterisation of cartilage intermediate layer protein (CILP)-induced arthropathy in mice, Annals of the rheumatic diseases 63(3) (2004) 252–8 10.1136/ard.2003.008045

[18] S. Seki, Y. Kawaguchi, K. Chiba, Y. Mikami, H. Kizawa, T. Oya, F. Mio, M. Mori, Y. Miyamoto, I. Masuda, T. Tsunoda, M. Kamata, T. Kubo, Y. Toyama, T. Kimura, Y. Nakamura, S. Ikegawa, A functional SNP in CILP, encoding cartilage intermediate layer protein, is associated with susceptibility to lumbar disc disease, Nat Genet 37(6) (2005) 607–12 10.1038/ng1557

[19] A.R. Armiento, M. Alini, M.J. Stoddart, Articular fibrocartilage - Why does hyaline cartilage fail to repair?, Adv Drug Deliv Rev 146 (2019) 289–305 10.1016/j.addr.2018.12.015

[20] M. Tschaikowsky, S. Brander, V. Barth, R. Thomann, B. Rolauffs, B.N. Balzer, T. Hugel, The articular cartilage surface is impaired by a loss of thick collagen fibers and formation of type I collagen in early osteoarthritis, Acta Biomater 146 (2022) 274–283 10.1016/j.actbio.2022.04.036

[21] B. Kwok, P. Chandrasekaran, C. Wang, L. He, R.L. Mauck, N.A. Dyment, E. Koyama, L. Han, Rapid specialization and stiffening of the primitive matrix in developing articular cartilage and meniscus, Acta Biomater 168 (2023) 235–251 10.1016/j.actbio.2023.06.047

[22] S. Gross, T. Thum, TGF-beta Inhibitor CILP as a Novel Biomarker for Cardiac Fibrosis, JACC Basic Transl Sci 5(5) (2020) 444–446 10.1016/j.jacbts.2020.03.013

[23] H. Li, X. Jiang, Y. Xiao, Y. Zhang, W. Zhang, M. Doherty, J. Nestor, C. Li, J. Ye, T. Sha, H. Lyu, J. Wei, C. Zeng, G. Lei, Combining single-cell RNA sequencing and population-based studies reveals hand osteoarthritis-associated chondrocyte subpopulations and pathways, Bone Res 11(1) (2023) 58 10.1038/s41413-023-00292-7

[24] X. Jiang, B.R. Stockwell, M. Conrad, Ferroptosis: mechanisms, biology and role in disease, Nat Rev Mol Cell Biol 22(4) (2021) 266–282 10.1038/s41580-020-00324-8

[25] D. Tang, X. Chen, R. Kang, G. Kroemer, Ferroptosis: molecular mechanisms and health implications, Cell research 31(2) (2021) 107–125 10.1038/s41422-020-00441-1

[26] Z. Hu, L. Chen, J. Zhao, W. Zhang, Z. Jin, Y. Sun, Z. Li, B. Chang, P. Shen, Y. Yang, Lipoxin A(4) ameliorates knee osteoarthritis progression in rats by antagonizing ferroptosis through activation of the ESR2/LPAR3/Nrf2 axis in synovial fibroblast-like synoviocytes, Redox Biol 73 (2024) 103143 10.1016/j.redox.2024.103143

[27] X. Sun, Z. Ou, R. Chen, X. Niu, D. Chen, R. Kang, D. Tang, Activation of the p62-Keap1-NRF2 pathway protects against ferroptosis in hepatocellular carcinoma cells, Hepatology 63(1) (2016) 173–84 10.1002/hep.28251

[28] L. Liu, J. Du, S. Yang, B. Zheng, J. Shen, J. Huang, L. Cao, S. Huang, X. Liu, L. Guo, C. Li, C. Ke, X. Peng, D. Guo, H. Peng, SARS-CoV-2 ORF3a sensitizes cells to ferroptosis via Keap1-NRF2 axis, Redox Biol 63 (2023) 102752 10.1016/j.redox.2023.102752

[29] Y. Yan, A. Lu, Y. Dou, Z. Zhang, X.Y. Wang, L. Zhai, L.Y. Ai, M.Z. Du, L.X. Jiang, Y.J. Zhu, Y.J. Shi, X.Y. Liu, D. Jiang, J.C. Wang, Nanomedicines Reprogram Synovial Macrophages by Scavenging Nitric Oxide and Silencing CA9 in Progressive Osteoarthritis, Advanced science (Weinheim, Baden-Wurttemberg, Germany) 10(11) (2023) e2207490 10.1002/advs.202207490

[30] F. Maier, C.G. Lewis, D.M. Pierce, The evolving large-strain shear responses of progressively osteoarthritic human cartilage, Osteoarthritis Cartilage 27(5) (2019) 810–822 10.1016/j.joca.2018.12.025

[31] K.P.H. Pritzker, S. Gay, S.A. Jimenez, K. Ostergaard, J.P. Pelletier, P.A. Revell, D. Salter, W.B. van den Berg, Osteoarthritis cartilage histopathology: grading and staging, Osteoarthritis and Cartilage 14(1) (2006) 13–29 10.1016/j.joca.2005.07.014

[32] M. Mazor, T.M. Best, A. Cesaro, E. Lespessailles, H. Toumi, Osteoarthritis biomarker responses and cartilage adaptation to exercise: A review of animal and human models, Scand J Med Sci Sports 29(8) (2019) 1072–1082 10.1111/sms.13435

[33] X. Wang, Q. Wu, R. Zhang, Z. Fan, W. Li, R. Mao, Z. Du, X. Yao, Y. Ma, Y. Yan, W. Sun, H. Wu, W. Wei, Y. Hu, Y. Hong, H. Hu, Y.W. Koh, W. Duan, X. Chen, H. Ouyang, Stage-specific and location-specific cartilage calcification in osteoarthritis development, Annals of the rheumatic diseases 82(3) (2023) 393–402 10.1136/ard-2022-222944

[34] J.T. Makela, Z.S. Rezaeian, S. Mikkonen, R. Madden, S.K. Han, J.S. Jurvelin, W. Herzog, R.K. Korhonen, Site-dependent changes in structure and function of lapine articular cartilage 4 weeks after anterior cruciate ligament transection, Osteoarthritis Cartilage 22(6) (2014) 869–78 10.1016/j.joca.2014.04.010

[35] T. Li, J. Liu, M. Guo, F.C. Bin, Q. Duan, X.Z. Dong, F. Jin, K. Fujita, M.L. Zheng, Femtosecond Laser Maskless Optical Projection Lithography of Cartilage PCM Inspired 3D Protein Matrix to Chondrocyte Phenotype, Adv Healthc Mater (2024) e2400849 10.1002/adhm.202400849

[36] Y. Ling, W. Zhang, P. Wang, W. Xie, W. Yang, D.A. Wang, C. Fan, Three-dimensional (3D) hydrogel serves as a platform to identify potential markers of chondrocyte dedifferentiation by combining RNA sequencing, Bioactive materials 6(9) (2021) 2914–2926 10.1016/j.bioactmat.2021.02.018

[37] R. Ossendorff, S. Grad, T. Tertel, D.C. Wirtz, B. Giebel, V. Börger, F.A. Schildberg, Immunomodulatory potential of mesenchymal stromal cell-derived extracellular vesicles in chondrocyte inflammation, Frontiers in immunology 14 (2023) 1198198 10.3389/fimmu.2023.1198198

[38] H. Kwon, W.E. Brown, S.A. O’Leary, J.C. Hu, K.A. Athanasiou, Rejuvenation of extensively passaged human chondrocytes to engineer functional articular cartilage, Biofabrication 13(3) (2021) 10.1088/1758-5090/abd9d9

[39] Y. Zhang, Z. Wang, C. Zong, X. Gu, S. Fan, L. Xu, B. Cai, S. Lu, Platelet-rich plasma attenuates the severity of joint capsule fibrosis following post-traumatic joint contracture in rats, Front Bioeng Biotechnol 10 (2022) 1078527 10.3389/fbioe.2022.1078527

[40] R.S. Watson, E. Gouze, P.P. Levings, M.L. Bush, J.D. Kay, M.S. Jorgensen, E.A. Dacanay, J.W. Reith, T.W. Wright, S.C. Ghivizzani, Gene delivery of TGF-β1 induces arthrofibrosis and chondrometaplasia of synovium in vivo, Laboratory investigation; a journal of technical methods and pathology 90(11) (2010) 1615–27 10.1038/labinvest.2010.145

[41] L. Weng, J. Ye, F. Yang, S. Jia, M. Leng, B. Jia, C. Xu, Y. Zhao, R. Liu, Y. Xiong, Y. Zhou, J. Zhao, M. Zheng, TGF-β1/SMAD3 Regulates Programmed Cell Death 5 That Suppresses Cardiac Fibrosis Post-Myocardial Infarction by Inhibiting HDAC3, Circulation research 133(3) (2023) 237–251 10.1161/circresaha.123.322596

[42] Y. Miao, Y. Chen, F. Xue, K. Liu, B. Zhu, J. Gao, J. Yin, C. Zhang, G. Li, Contribution of ferroptosis and GPX4’s dual functions to osteoarthritis progression, EBioMedicine 76 (2022) 103847 10.1016/j.ebiom.2022.103847

[43] L. Yu, S. Cavelier, B. Hannon, M. Wei, Recent development in multizonal scaffolds for osteochondral regeneration, Bioactive materials 25 (2023) 122–159 10.1016/j.bioactmat.2023.01.012

[44] A.R. Armiento, M.J. Stoddart, M. Alini, D. Eglin, Biomaterials for articular cartilage tissue engineering: Learning from biology, Acta Biomater 65 (2018) 1–20 10.1016/j.actbio.2017.11.021

[45] H. Zhang, L. Ji, Y. Yang, Y. Wei, X. Zhang, Y. Gang, J. Lu, L. Bai, IF3.2The Therapeutic Effects of Treadmill Exercise on Osteoarthritis in Rats by Inhibiting the HDAC3/NF-KappaB Pathway in vivo and in vitro, Front Physiol 10 (2019) 1060 10.3389/fphys.2019.01060

[46] K. Richter, A. Konzack, T. Pihlajaniemi, R. Heljasvaara, T. Kietzmann, Redox-fibrosis: Impact of TGFβ1 on ROS generators, mediators and functional consequences, Redox Biol 6 (2015) 344–352 10.1016/j.redox.2015.08.015

[47] Q. Ru, Y. Li, W. Xie, Y. Ding, L. Chen, G. Xu, Y. Wu, F. Wang, Fighting age-related orthopedic diseases: focusing on ferroptosis, Bone Res 11(1) (2023) 12 10.1038/s41413-023-00247-y

[48] B.R. Stockwell, Ferroptosis turns 10: Emerging mechanisms, physiological functions, and therapeutic applications, Cell 185(14) (2022) 2401–2421 10.1016/j.cell.2022.06.003

[49] H. Zhang, J. Pan, S. Huang, X. Chen, A.C.Y. Chang, C. Wang, J. Zhang, H. Zhang, Hydrogen sulfide protects cardiomyocytes from doxorubicin-induced ferroptosis through the SLC7A11/GSH/GPx4 pathway by Keap1 S-sulfhydration and Nrf2 activation, Redox Biol 70 (2024) 103066 10.1016/j.redox.2024.103066

[50] T. Lin, S. Peng, S. Lu, S. Fu, D. Zeng, J. Li, T. Chen, T. Fan, C. Lang, S. Feng, J. Ma, C. Zhao, B. Antony, F. Cicuttini, X. Quan, Z. Zhu, C. Ding, Prediction of knee pain improvement over two years for knee osteoarthritis using a dynamic nomogram based on MRI-derived radiomics: a proof-of-concept study, Osteoarthritis and cartilage 31(2) (2023) 267–278 10.1016/j.joca.2022.10.014

[51] T. Cui, R. Liu, Y. Jing, J. Fu, J. Chen, Development of machine learning models aiming at knee osteoarthritis diagnosing: an MRI radiomics analysis, J Orthop Surg Res 18(1) (2023) 375 10.1186/s13018-023-03837-y

[52] T.E. McAlindon, R.R. Bannuru, M.C. Sullivan, N.K. Arden, F. Berenbaum, S.M. Bierma-Zeinstra, G.A. Hawker, Y. Henrotin, D.J. Hunter, H. Kawaguchi, K. Kwoh, S. Lohmander, F. Rannou, E.M. Roos, M. Underwood, OARSI guidelines for the non-surgical management of knee osteoarthritis, Osteoarthritis and cartilage 22(3) (2014) 363–88 10.1016/j.joca.2014.01.003

[53] S.S. Glasson, T.J. Blanchet, E.A. Morris, The surgical destabilization of the medial meniscus (DMM) model of osteoarthritis in the 129/SvEv mouse, Osteoarthritis and cartilage 15(9) (2007) 1061–9 10.1016/j.joca.2007.03.006

[54] Y. Yang, Y. Wang, Y. Kong, X. Zhang, H. Zhang, Y. Gang, L. Bai, The therapeutic effects of lipoxin A4 during treadmill exercise on monosodium iodoacetate-induced osteoarthritis in rats, Mol Immunol 103 (2018) 35–45 10.1016/j.molimm.2018.08.027

[55] Y. Yang, Y. Wang, Y. Kong, X. Zhang, H. Zhang, X. Feng, Z. Wang, P. Gao, M. Yan, L. Bai, F. Li, Moderate Mechanical Stimulation Protects Rats against Osteoarthritis through the Regulation of TRAIL via the NF-kappaB/NLRP3 Pathway, Oxid Med Cell Longev 2020 (2020) 6196398 10.1155/2020/6196398

[56] H. Zhang, L. Ji, Y. Yang, Y. Wei, X. Zhang, Y. Gang, J. Lu, L. Bai, The Therapeutic Effects of Treadmill Exercise on Osteoarthritis in Rats by Inhibiting the HDAC3/NF-KappaB Pathway in vivo and in vitro, Front Physiol 10 (2019) 1060 10.3389/fphys.2019.01060

[57] Q.F. Wills, K.J. Livak, A.J. Tipping, T. Enver, A.J. Goldson, D.W. Sexton, C. Holmes, Single-cell gene expression analysis reveals genetic associations masked in whole-tissue experiments, Nature biotechnology 31(8) (2013) 748–52 10.1038/nbt.2642

[58] E.J. Duncavage, M.C. Schroeder, M. O’Laughlin, R. Wilson, S. MacMillan, A. Bohannon, S. Kruchowski, J. Garza, F. Du, A.E.O. Hughes, J. Robinson, E. Hughes, S.E. Heath, J.D. Baty, J. Neidich, M.J. Christopher, M.A. Jacoby, G.L. Uy, R.S. Fulton, C.A. Miller, J.E. Payton, D.C. Link, M.J. Walter, P. Westervelt, J.F. DiPersio, T.J. Ley, D.H. Spencer, Genome Sequencing as an Alternative to Cytogenetic Analysis in Myeloid Cancers, The New England journal of medicine 384(10) (2021) 924–935 10.1056/NEJMoa2024534

[59] S. Tasaki, J. Xu, D.R. Avey, L. Johnson, V.A. Petyuk, R.J. Dawe, D.A. Bennett, Y. Wang, C. Gaiteri, Inferring protein expression changes from mRNA in Alzheimer’s dementia using deep neural networks, Nature communications 13(1) (2022) 655 10.1038/s41467-022-28280-1

[60] N. Gerwin, A.M. Bendele, S. Glasson, C.S. Carlson, The OARSI histopathology initiative - recommendations for histological assessments of osteoarthritis in the rat, Osteoarthritis and cartilage 18 Suppl 3 (2010) S24–34 10.1016/j.joca.2010.05.030

[61] E. Charlier, C. Deroyer, F. Ciregia, O. Malaise, S. Neuville, Z. Plener, M. Malaise, D. de Seny, Chondrocyte dedifferentiation and osteoarthritis (OA), Biochem Pharmacol 165 (2019) 49-65 10.1016/j.bcp.2019.02.036

[62] Y. Huang, Y. Liu, Q. Huang, S. Sun, Z. Ji, L. Huang, Z. Li, X. Huang, W. Deng, T. Li, TMT-Based Quantitative Proteomics Analysis of Synovial Fluid-Derived Exosomes in Inflammatory Arthritis, Frontiers in Immunology 13 (2022) 10.3389/fimmu.2022.800902

[63] J.G. Meyer, N.M. Niemi, D.J. Pagliarini, J.J. Coon, Quantitative shotgun proteome analysis by direct infusion, Nature methods 17(12) (2020) 1222–1228 10.1038/s41592-020-00999-z

